# Binding free-energy landscapes of small molecule binder and non-binder to FMN riboswitch: All-atom molecular dynamics

**DOI:** 10.1101/2023.07.01.547313

**Authors:** Junichi Higo, Gert-Jan Bekker, Narutoshi Kamiya, Ikuo Fukuda, Yoshifumi Fukunishi

**Author notes:** Correspondence and requests for materials should be addressed to J.H.

## Abstract

Binding of a small and flexible molecule, ribocil A (non-binder) or B (binder), to the deep pocket of the aptamer domain of the FMN riboswitch was studied by mD-VcMD, which is a generalized-ensemble method based on molecular dynamics (MD) simulation. Ribocil A and B are structurally similar because they are optical isomers mutually. In the initial conformation of simulation, both ligands and the aptamer were completely dissociated in explicit solvent. The resultant free-energy landscape of ribocil B binding to the aptamer was funnel-like, whereas that of ribocil A was rugged, which agrees qualitatively with an experiment. When entering the gate (named “front gate”) of the pocket, the ligand interacted with the aptamer by native and non-native π-π stackings, and the stackings restrained the molecular orientation of the ligands to be advantageous to reach the binding site smoothly. The simulation showed another pathway, which also led the ligands to the binding site. Its gate (maned “rear gate”) located completely opposite to the front gate on the aptamer’s surface. However, approach from the rear gate required overcoming a free-energy barrier before reaching the binding site, and the ligands should rotate largely and sharply at the free-energy barrier. This ligand’s orientation property is discussed referring to a ligand orientation selection mechanism exserted by a membrane protein capturing its ligand.

## 1. Introduction

Computational drug discovery of RNA has been one of emerging topics (Bernetti et al., 2022; Manigrasso et al., 2021; Morishita, 2023; Childs-Disney et al., 2022; Bagnolini et al., 2023; Kognole et al., 2022) and riboswitches are popular targets. Riboswitches are located in the 5’ or 3’ untranslated regions of mRNA and regulate translations of mRNAs when small molecules bind to riboswitches (Nahvi et al., 2002; Mironov et al., 2002; Winkler et al., 2002a; Winkler et al., 2002b), as shown in functioning of allosteric proteins. The riboswitch consists of two regions: Aptamer domain and expression platform. Upon binding of a small ligand to the binding site of the aptamer domain, the structure of the expression platform varies, and then a gene–expression process is switched on or off by the effect of the structural variation. In general, the ligand-binding rate to a receptor (the riboswitch in this study) depends on the ligand concentration around the receptor. Therefore, the riboswitch has a role of switching on or off for the translation depending on the ligand concentration (Serganov & Nudler, 2013). Because riboswitches have been found in pathogenic bacteria, a compound, which inhibits binding of the genuine ligand binding to the riboswitch, can be a drug candidate (Warner et al., 2018).

Riboflavin (also known as vitamin B_2_) is metabolized and flavin mononucleotide (FMN) is yielded in cell. The FMN riboswitch (Mironov et al., 2002; Winkler et al., 2002b), or known as RFN element (Gelfand et al., 1999), is a riboswitch of prokaryote and the aptamer domain binds FMN. This binding results in suppression of riboflavin’s biosynthesis with inducing a structural change in its expression platform (Winkler et al., 2002a; Winkler et al., 2002b). When the concentration of FMN raised in cell, the probability of complex formation of FMN and the aptamer domain raises. Thus, the FMN riboswitch works as a sensor for feed-back control of the biosynthesis of riboflavin (Gelfand et al., 1999; Vitreschak, et al., 2002).

The structure of the aptamer domain of the *Fusobacterium nucleatum* FMN riboswitch has been studied experimentally (Vicens, 2011; Serganov et al., 2009; Rizvi et al., 2018; Vicens et al., 2018). The complex structure of ribocil B and the aptamer domain was solved by an X-ray crystallography (PDB entry = 5c45; resolution 2.93 Å) (Howe et al., 2015). Figure S1 of supplementary Information (SI) illustrates the ligand bound to the deep pocket of the aptamer domain. Portions of the aptamer domain are referred to as P1–P6 (Wilt et al., 2020) as shown in Figure S1a–c. An experiment has studied binding of some compounds to the aptamer domain and reported that ribocil B binds to the aptamer domain more strongly than ribocil A does, although ribocil A and B are very similar compounds (optical isomers) to each other (Figures 1a and b) (Howe, et al., 2016). Interestingly, the binding modes of ribocil A and B to the aptamer domain are almost identical (Howe, et al., 2016), whereas the X-ray complex structure was nor reported for ribocil A.

**Figure 1.**
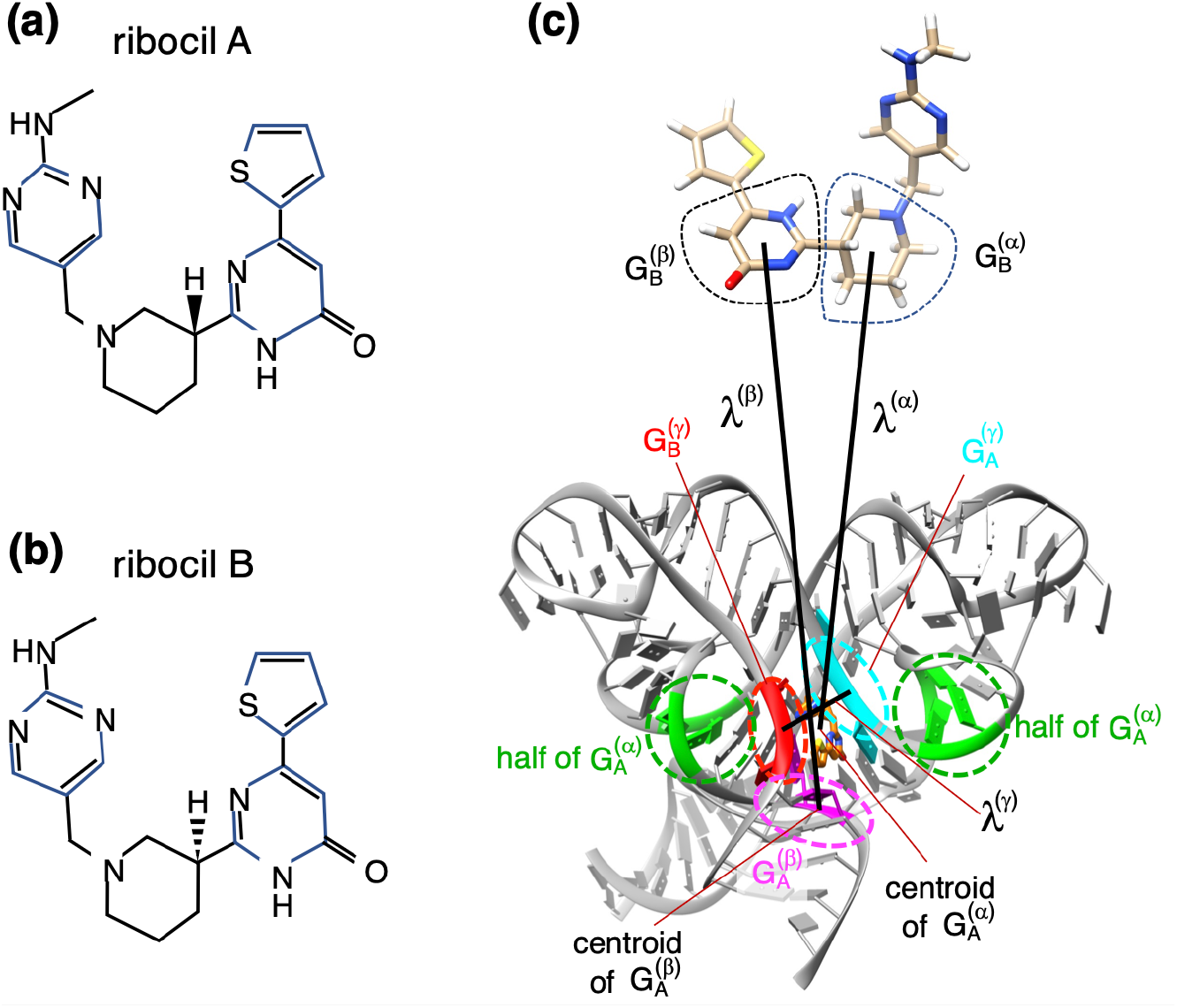
Examined small molecules for mD-VcMD simulations. (a) Ribocil A (non-binder) and (b) ribocil B (binder), which are optical isomers mutually. Ligands are flexible in simulation because all the dihedral bonds are rotatable. (c) Definition of three RCs: *λ*^(*α*)^, *λ*^(*β*)^, and *λ*^(*γ*)^. Method to set an RC (*λ*^(*h*)^; *h = α, β*, or *γ*) with using two atom groups 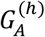 and 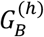 is explained in the main text and Section 2 of SI. See Table S1 of SI for details of atom groups adopted actually in the present study. Atom groups 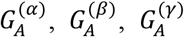 and 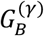 are set in the aptamer domain, and 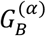 and 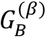 are done in ribocil. The atom group 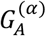 presented in green is split into two regions of the aptamer domain.

Figure S1 defines three directions to view the aptamer domain of the FNM riboswitch: “front view” (Figures S1a and d), “side view” (Figures S1b and e), and “rear view” (Figures S1c and f). The aptamer domain is presented by the front view in many papers. Interestingly, the ligand ribocil B is visible in both the front and rear views (the magenta molecule in Figures S1d and f). Figure S2 displays the apo and holo forms of the aptamer domain, where ribicil B was removed in the apo form (Figure S2b). This figure indicates that the binding pocket forms a tunnel between the front and rear sides of the aptamer domain in both forms. Existence of tunnel suggests that the ligand may reach the binding position from both the front and rear sides.

A molecular dynamics simulation can provide dynamic processes occurring in a biomolecular system. Especially, enhanced sampling (generalized-ensemble methods) reproduces rare and large conformational motions in the molecular system (Higo et al., 2012; Kasahara et al., 2019; Fukunishi et al., 2022). The multi-dimensional virtual-system coupled molecular dynamics (mD-VcMD) (Higo et al., 2020a) is one of the enhanced sampling methods designed for investigating molecular binding process. To perform an mD-VcMD simulation, a user introduces a set of multiple reaction coordinates (RCs) in advance. Then, the method enhances the motions regarding the multiple RCs. Importantly, a statistical weight (thermodynamic weight) at a simulation temperature is assigned to all the sampled conformations (snapshots). Therefore, the ensemble of snapshots can be regarded a thermally equilibrated ensemble (canonical ensemble) of the snapshots. This method was applied to various ligand–receptor systems (Higo, et al., 2020b; Higo et al., 2021; Hayami et al., 2021; Higo et al., 2019; Higo et al., 2022; Xie et al., 2023). We have proposed three variants of mD-VcMD so far (Higo et al., 2020a): The original, a subzone-based, and genetic algorithm-guided mD-VcMD simulations.

In this paper we investigate computationally the mechanism of ribocil–aptamer complex formation focusing two issues: i) Whether ribocil B binds to the aptamer domain more strongly than ribocil A does and ii) whether ribocil approaches the binding pocket from both the front and rear side of the aptamer domain. For this purpose, we perform mD-VcMD starting the simulation from a conformation where ribocil is far from the aptamer domain. The simulation samples widely the conformational space and produce conformational ensembles for two systems: A system of the aptamer domain and ribocil B and a system of the aptamer domain and ribocil A. Both ensembles consist of unbound and bound (complex) conformations. Then, we calculate free-energy landscapes for the two systems, from which the two issues mentioned above are discussed.

## 2. Materials and Methods

In this work, the mD-VcMD simulation is performed to obtain the conformational ensemble of the system. The system consists of a receptor (the aptamer domain of the *F. nucleatum* FMF riboswitch) and a ligand (ribocil A or B), and the ensemble is used to obtain the free-energy landscape from the ensemble. When the ligand is ribocil A, the system is referred to as “Ribo-A” and when it is ribocil B, it is done to as “Ribo-B”.

For conformational sampling, we use the subzone-based mD-VcMD, which is one of the three variants of the mD-VcMD methods (Higo et al., 2020a), although the method is simply referred to as mD-VcMD in this paper. A statistical weight (i.e., thermodynamic weight at the simulation temperature) is assigned to any sampled conformation (i.e., snapshots). Then the ensemble of the weighted snapshots (i.e., canonical ensemble) is used to calculate various physical quantities at equilibrium.

Below, we first explain the simulation system. Next, we introduce multiple reaction coordinates (RCs), along which the conformational sampling is enhanced. Third, we explain briefly the mD-VcMD sampling method. Last, we explain procedures to calculate some physical quantities.

### 2.1. Molecular system and the initial conformation of simulation

The structure of the aptamer domain was taken from an X-ray crystallography (PDB entry = 6wjr; 2.7 Å resolution), which is an apo form (i.e., unbound form) of the aptamer domain (Wilt et al., 2020). The aptamer domain was put at the center of a periodic boundary box filled by solvent (97.9582 Å × 97.9582 Å × 97.9582 Å), and ribocil A or B was put at the corner of the box (see Figure S3a of SI). Then, an NPT simulation was applied to the system, and the resultant conformation was used for the initial conformation of the mD-VcMD simulation (Figure S3b). Details for generating the initial conformation are explained in Section 1 of SI. The number of atoms is 92,628, which is the same for the two systems: 3,572 RNA atoms, 49 ligand atoms, 29,602 water molecules, 15 Mg ions, 133 K ions and 53 Cl ions. Mg ions were located at the Mg-binding sites of the riboswitch (see Section 1 of SI for details). The other ions were introduced to set the ionic concentration to an experimental condition (0.1 M) with neutralizing the net charge of the entire systems (Howe et al., 2015). The resultant box size from the NPT simulation was 97.3511 Å × 97.3511 Å × 97.3511 Å and 97.3304 Å × 97.3304 Å × 97.3304 Å for the Ribo-A and Ribo-B systems, respectively.

In the initial conformation of mD-VcMD simulation, the ribocil (A or B) is near the corner of the periodic in both the Ribo-A and Ribo-B systems as shown in Figure S3b. Therefore, the ribocil–aptamer complex structure is not the initial conformation of simulation but a conformation to be searched by the simulation.

The force field for the aptamer domain was from the ff99bsc0χOL3 (or χ_OL3_) (Zgarbová et al., 2011) and those for water molecule were from the 3-point optimal point charge (OPC3) water model (Izadi & Onufriev, 2016). The force field parameters for Mg ion were from a study (Li et al., 2020) and those for K and Cl ions were from another study (Sengupta et al., 2021).

The chemical structures for ribocil A and B are shown in Figures 1a and b. Force field parameters for ribocil A and B were modeled by us. First, the atomic partial charges of ribocil A or B were calculated quantum-chemically using Gaussian09 (Frisch et al., 2009) at the HF/6-31G* level, followed by RESP fitting (Bayly et al., 1993). Then, the obtained atomic partial charges were incorporated into a general amber force field file 2 (GAFF2) (Wang et al., 2004), which was designed to be compatible with conventional AMBER force-fields.

### 2.2. Three RCs introduced to control system’s motions

The mD-VcMD simulation enhances the conformational sampling in a space constructed by multiple RCs: *λ*^(*h*)^ (*h = α, β, γ*, …). Thus, the RCs should be set beforehand. In this study, an RC, *λ*^(*h*)^, is defined by the inter-centroid distance between two atom groups, 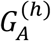 and 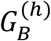, as shown in Section 2 of SI. Selection of RCs is important to increase the sampling efficiency, although RCs can be set arbitrarily in theory. We imposed three conditions on the RCs: Variations of RCs can control (i) ribocil approaching to (or departing from) the aptamer domain, (ii) ribocil’s orientation relative to the aptamer domain, and (iii) the gate opening (or closing) of the binding pocket of the aptamer domain. Although various RCs can satisfy the three conditions, we set the RCs in a straightforward manner as shown in Figure 1c for both the Ribo-A and Ribo-B systems. This is because ribocil A and ribocil B overlaps very well to each other in the complex structures (Howe, et al., 2016) as mentioned in the *Introduction* section.

The atom groups used for the present RCs (Figure 1c) are given in Section 3 and Table S1 of SI. By the introduction of the three RCs, “multi-dimensional RC (mD RC)” becomes “three-dimensional (3D RC)” actually. We briefly explain the roles of the RCs. Two atom groups 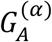 and 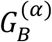 are introduced to define the first RC *λ*^(*α*)^, and increment or decrement of *λ*^(*α*)^ corresponds to ribocil approaching to or departing from the aptamer domain, respectively. This relates to condition (i). The atom groups 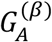 and 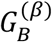 define *λ*^(*β*)^ and motions of *λ*^(*β*)^ also related to condition (i). However, if *λ*^(*α*)^ decreases and *λ*^(*β*)^ increases simultaneously, or if *λ*^(*α*)^ increases and *λ*^(*β*)^ decreases, then the ligand rotates: Condition (ii) is satisfied. Last, 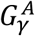 and 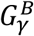 define the third RC *λ*^(*γ*)^, and increment (or decrement) of *λ*^(*γ*)^ induces opening (or closing) of the gate of the binding pocket: Condition (iii) is satisfied.

Similar RC setting was used in our previous studies (Higo et al., 2021; Hayami et al., 2021; Higo et al., 2019; Higo et al., 2022). The atom groups 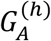 and 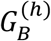 were set as independent of the chemical properties of the ligand and receptor. The atom groups were set conveniently to establish the three conditions (i), (ii) and (iii) explained above. One might infer that enhancement of the conformational motions in the 3D RC space affects the resultant conformational distribution and snapshots because a bias is introduced for the enhancement. This criticism is true for conventional MD simulation. In mD-VcMD, however, the bias effect is removed completely from the resultant ensemble using a reweighting technique (Higo et al., 2020a). The estimated thermodynamic weight assigned to a conformation is one without the bias effect in mD-VcMD.

### 2.3. mD-VcMD and a conformational ensemble

The mD-VcMD samples the conformation in a region of 3D RC space at 300 K, and the region should be defined in advance by user. For this purpose, we set the minimum and maximum boundaries (denoted as 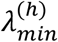 and 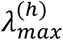, respectively) for each RC. Then, the conformation moves in the range of 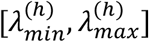. Importantly, the conformation should adopt both bound and unbound conformations in the range. The actual values of the boundaries are presented in Table S2 of SI. We set 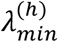 to 0.0Å for *h = α* and *β*. This value of zero is the smallest in theory. The 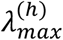 (83.1178 Å) for *h = α* and *β* is large enough to produce unbound conformations. Recall that *λ*^(*γ*)^ is the gate width of the binding pocket of the aptamer domain. The value of 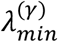 (5.0 Å) is small enough because it is likely that ribocil cannot not pass through the gate. The radius of a heavy atom is 2 Å approximately. When 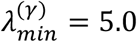 Å, only a space of 1 Å remains, which is smaller than the diameter of a heavy atom (5 Å *=* 2 Å + 2 Å + 1 Å). The value of 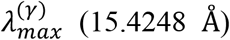, which corresponds to the maximum gate width, allowed ribocil to pass through the gate as shown later.

Next, we divided the *λ*^(*h*)^ axis into small 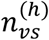 zones: Table S2 of SI lists the actual value of 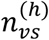. The lower and upper boundaries for the *i*-th zone along *λ*^(*h*)^ are denoted as 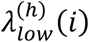 and 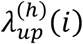, respectively, whose values are listed in Table S3. We refer to a zone along an RC axis as a 1D zone. A relation of 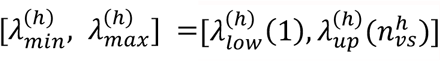 is satisfied because the 1D zones were introduced by dividing the range of 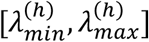 into 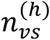 zones. In the 3D RC space, a zone is defined three-dimensionally: A set of *i*-th, *j*-th, and *k*-th 1D zones along the *λ*^(*α*)^-, *λ*^(*β*)^- and *λ*^(*γ*)^-axis, respectively, corresponds to a “3D zone” whose index is (*i, j, k*). The ranges for this 3D zone are 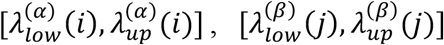 and 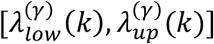 along the *λ*^(*α*)^-, *λ*^(*β*)^-, and *λ*^(*γ*)^-axis, respectively.

Assume that the conformation of the system is currently in the 3D zone (*i, j, k*) at time *= t*. We refer to this 3D zone as the “current zone”. In mD-VcMD, the conformation is confined in the current zone for a short interval, Δτ_*trns*_, of simulation: A restoring force towards the current zone is applied to the system only when the conformation is flying to outside the current zone, and the conformation inside the current zone moves without the restoring force (Higo et al., 2020a). Then, at a time passage of Δτ_*trns*_ (time *= i* + Δτ_*trns*_), the conformation may transition from (*i, j, k*) to another zone (*i*′, *j*′, *k*′), which is one of the zones adjacent to (*i, j, k*), with using a transition matrix *B* (Higo et al., 2020a). Now, the current zone is updated to (*i*′, *j*′, *k*′). A matrix element *B*_(*i,j,k*)→(*i*′, *j*′, *k*′)_ is the transition probability from zone (*i, j, k*) to zone (*i*′, *j*′, *k*′). In the present study we set Δτ_*trns*_ *=* 20 *ps*, which corresponds to 1 × 10^4^ simulation steps (a time step of simulation = 2 *fs*).

Repeating this procedure, the conformation of the system moves in the wide 3D RC space. However, because the conformational space is too large to be sampled in a single simulation, the mD-VcMD simulation is performed iteratively. When iteration *M* has finished, a quantity 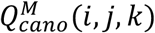 is calculated from iterations performed so far (i.e., iterations 1–*M*): 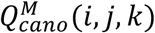 is a probability distribution of the system existing in zone (*i, j, k*) at equilibrium (Higo et al., 2020a). In other words, 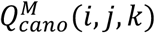 is a system’s canonical distribution (i.e., a thermally equilibrated distribution at the simulation temperature *T*) in a 3D zone (*i, j, k*) when convergence of 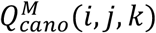 is obtained in iterations from 1 to *M*. The transition matrix *B* for the (*M* + 1)-th iteration is calculated from 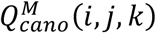 whose protocol is given in our earlier studies (Higo et al., 2020a; 2020b; Higo et al., 2021; Hayami et al., 2021; Higo et al., 2019; Higo et al., 2022). Then the (*M* + 1)-th iteration is started. The transition matrix for the first iteration is set exceptionally as: *B*_(*i,j,k*)→(*i′, j′, k′*)_ *=* 1.0, which means that the transition from the current zone to any adjacent zones occurs equally.

The simulation is terminated when a convergence criterion is satisfied for 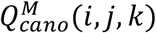 in iteration *M*, and then 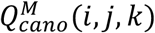 is expressed simply as 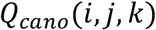. The convergence is discussed in the *Results and Discussion* section. A snapshot of the system is stored once every 1 × 10^5^ steps (0.2 ns) in all executed iterations. After convergence, a thermodynamic weight at simulation temperature *T* is assigned to each snapshot stored with using the probability distribution *Q*_*cano*_(*i, j, k*) (Higo et al., 2020a). Therefore, the snapshots construct a thermally equilibrated conformational ensemble (canonical ensemble) at *T* (300 K).

To increase further the sampling efficiency, we performed *N*_*run*_ (= 1,024) runs of the mD-VcMD simulation in parallel for each iteration, where the runs started from different conformations. Conformations from the *N*_*run*_ trajectories construct a canonical ensemble (Higo et al., 2009; Ikebe et al., 2011). Denoting the length of a single run as *L*, if the runs are initiated from various conformations, then the ensemble consisting of *N*_*run*_ trajectories covers a conformational space more widely than a long single run, whose length is *N*_*run*_ × *L*, does (Higo et al., 2009; Ikebe et al., 2011). If the *M*-th iteration has finished, we have snapshots sampled from iterations 1 to *M*. Then, the initial conformations of the *N*_*run*_ runs for the (*M* + 1)-th iteration were selected from those snapshots so that they are distributed as evenly as possible in the 3D RC space. Exceptionally, all the runs for the first iteration start from a single conformation obtained from the NPT simulation (Figure S3b).

We set the simulation length of each of *N*_*run*_ runs to 1 × 10^6^ steps (1 × 10^6^ × 2 fs *=* 2.0 ns) for both Ribo-A and Ribo-B systems. Consequently, the total length for an iteration was 2.048 μs (=2.0 ns × 1,024). We performed 45 iterations for the present study, which corresponds to 92.16 μs (2.048 μs × 45) in all for each system. As mentioned above, a snapshot was stored every 1 × 10^5^ steps (0.2 ns) in each run. Therefore, 460,800 (92.16 μs/0.2 ns) snapshots were stored for each system. We calculated the distribution functions of various quantities at the simulation temperature (*T =* 300 K) from the resultant ensemble of snapshots.

The mD-VcMD algorithm was first implemented to a MD simulation program omegagene/myPresto (Kasahara et al., 2019) and next to GROMACS (Kutzner et al., 2019) with the PLUMED plug-in (Bonomi et al., 2019), which was also used for the current study. The simulation conditions were the following: LINCS (Hess et al., 1997) to constrain the covalent-bond lengths related to hydrogen atoms, the Nosé–Hoover thermostat (Nosé, 1984; Hoover, 1985) to control the simulation temperature, the zero-dipole summation method (Kamiya, et al., 2013; Fukuda, et al., 2012; Fukuda, et al., 2011) to compute the long-range electrostatic interactions accurately and quickly, a time-step of 2 fs (Δ*i =* 2 fs), and simulation temperature of 300 K. All simulations were performed on the TSUBAME3.0 supercomputer at the Tokyo Institute of Technology using GP-GPU. The force field parameters used for the present study were explained in the above subsection.

### 2.4. Some physical quantities to study binding mechanism

The mD-VcMD simulation produces a canonical ensemble of conformations (snapshots) at 300 K. This method assigns a thermodynamic weight at 300 K to any snapshot obtained from the sampling (Higo et al., 2020a). Therefore, using this ensemble, one can calculate a spatial density of the ligand’s centroid around the receptor at 300 K using the ensemble: See Section 4 of SI for details.

Here we briefly explain the notations used to express the spatial density: See Subsection 4.1 of SI for details. The 3D real space (not the 3D-RC space) is divided into cubes (*L*_*c*_ × *L*_*c*_ × *L*_*c*_), where *L*_*c*_ is the length of sides of the cube, and the center of a cube is denoted as ***r***_*cube*_. *L*_*c*_ is usually set to 2 Å and rarely to 0.5 Å. If no explanation is mentioned, it means that *L*_*c*_ is 2 Å in this paper. The spatial density *ρ*_*CM*_(***r***_*cube*_) is the density of the ligand’s centroid ***r***_*CM*_ in the cube centered at ***r***_*cube*_.

Next, we defined two molecular orientation vectors ***e***_↑_ and ***e***_←_ for ribocil (Subsection 4.2 of SI). The ***e***_↑_ is a unit vector parallel to a vector *ν*_↑_, which is defined from the centroid of Ring 2 and Ring 3 to that of Ring 1 and Ring 4. The other vector ***e***_←_ is a unit vector parallel to a vector *ν*_←_, which is defined from the centroid of Ring 3 to that of Ring 2. Figure S5 of SI shows visually ***e***_↑_ and ***e***_←_. In general, an expectation value of a quantity Ω in the cube centered at ***r***_*cube*_ is denoted as < Ω(***r***_*cube*_) >. Expression for < Ω(***r***_*cube*_) > is given in Equation S4 of SI. Then, setting as Ω *=* ***e***_←_ or ***e***_↑_, the vector fields for the ribocil’s molecular orientation, < ***e***_↑_(***r***_*cube*_) > and < ***e***_←_(***r***_*cube*_) >, are calculated.

The native complex structure (PDB: 5c45) shows three π-π stacking (native stacking) between aptamer’s bases and ribocil’s rings (see Table S4 of SI). To analyze the stacking, we introduced a quantitative procedure to assign the stacking to snapshots: See Subsection 4.3 of SI. We checked the snapshots and found eight of stacking patterns (see also Table S4), three of which were the native stackings and the other five were non-native stacking. Figures S7a–c of SI illustrate snapshots with native stacking, and Figures S7d–f of SI do those with non-native stacking. We introduced two quantities: 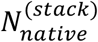 and 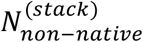, which are respectively the numbers of native and non-native stacking patterns in a snapshot.

## 3. Results

We performed 45 iterations of mD-VcMD and saved 460,800 snapshots as mentioned in the *Methods and Materials* section. In the *Results* section, we hirst show a distribution of the system in the 3D RC space that covered both the unbound and bound conformations. Next, we show that robocil B at the highest-density spot (the lowest free-energy position) was in the deep pocket of the aptamer domain, whereas that of ribocil A was not. We then analyze the binding mechanism from the conformational ensemble.

### 3.1. Conformational distribution in 3D-RC space

Figure 2 illustrates the computed *Q*_*cano*_(*i, j, k*), which is the equilibrated distribution of the system at 300 K in the 3D RC space (not in the 3D real space). As explained in the *Materials and Methods* section, the three indices *i, j* and *k* specify the position of a small zone in the 3D RC space. Because *Q*_*cano*_(*i, j, k*) is a probability at the zone, we normalized it as:

**Figure 2.**
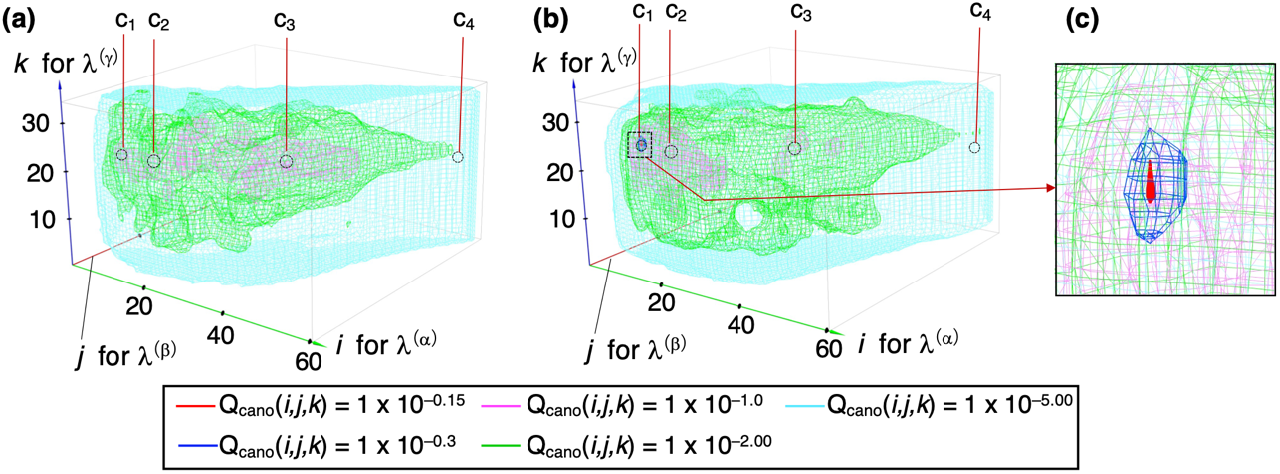
System’s canonical distribution 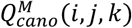 presented in the 3D RC space. 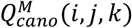 is computed from the 1’st to *M*-th iterations (*M* ≤ 45). Panels (a) and (b) are, respectively, distributions for Ribo-A and Ribo-B systems. (c) The close-up of the highest density region of panel (b). Three coordinate axes are indices *i, j*, and *k*, which specify a 3D zone. The position of the zone in the 3D RC space is: 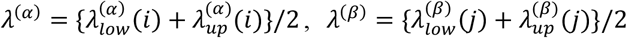 and 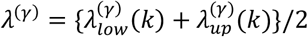. See Table S3 of SI for values of 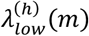 and 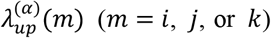. Snapshots from four sites *c*_1_– *c*_4_ of panels (a) and (b) are shown in Figure 3.

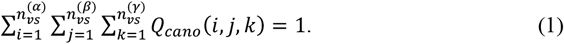

This normalization is important to compare *Q*_*cano*_ (*i, j, k*) between the Ribo-A and Ribo-B systems.

Figures 2b and c demonstrate that the Ribo-B system has a remarkable high-density spot (site *c*_1_ in Figure 2b) in the 3D RC space. In contrast, no such a remarkable high-density spot was found in the Ribo-A system (Figure 2a) with the unified density normalization (Equation 1). Therefore, the free-energy landscape for the Ribo-B system is funnel-like in the 3D RC space, while that for the Bibo-A system is rugged.

Conformations taken from sites *c*_1_ – *c*_4_ in Figure 2 are displayed in Figure 3a–d for the Ribo-A system and in Figure 3e–h for the Ribo-B system. Note that ribocil from *c*_1_ is in the binding pocket of the aptamer domain for both systems (Figures 3a and 3e). This is because conformations from *c*_1_ had similar RC values between the two systems (Figure 2). Because a high density was not assigned to *c*_1_ for the Ribo-A system, conformations from *c*_1_ in the Ribo-A system were thermodynamically less stable than those in the Ribo-B system. In the next subsection, we analyze details of the conformations in the binding pocket.

**Figure 3.**
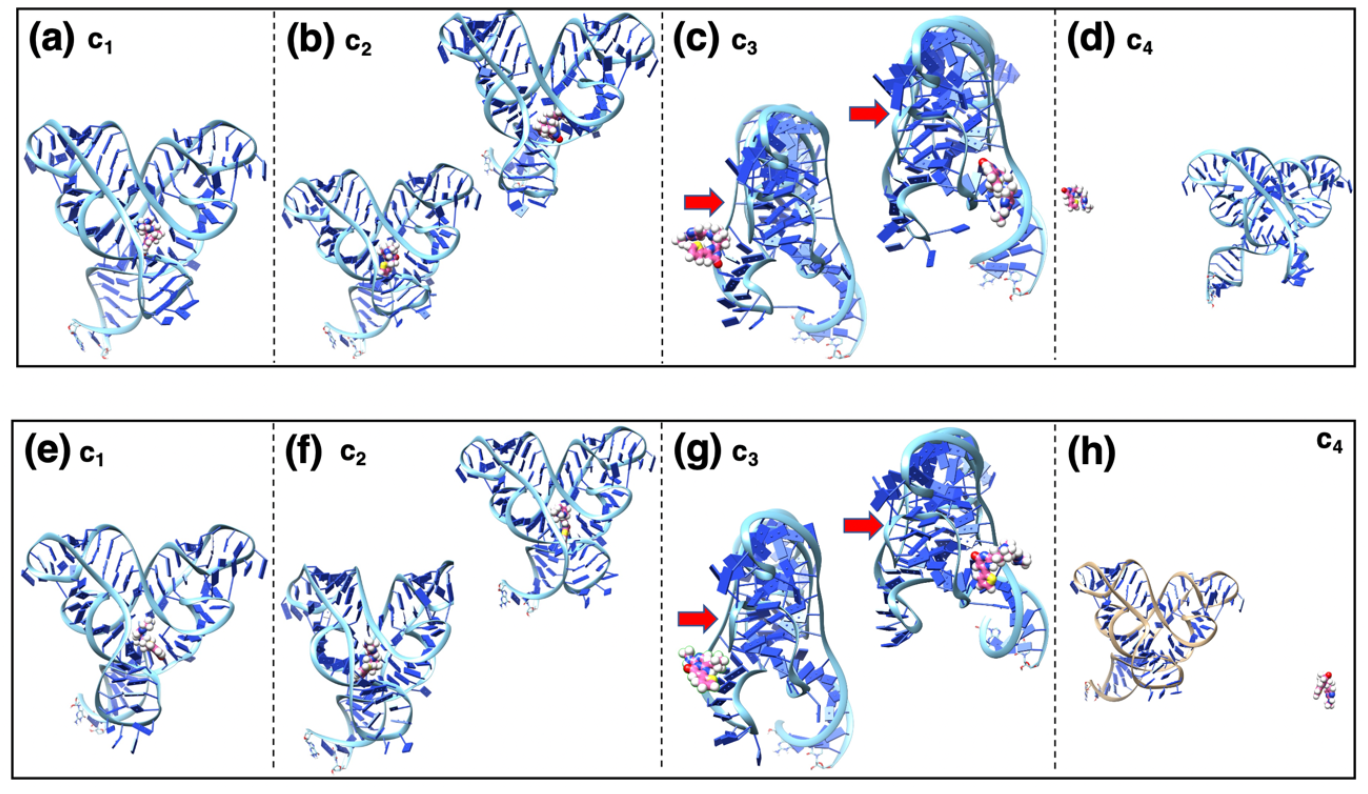
Conformations picked from sites *c*_*i*_ (*i =* 1, …, 4) whose positions in the 3D RC space are shown in Figure 2. Ribocils and aptamer are presented in magenta and light cyan, respectively. Panels (a), (b), (c) and (d) display snapshots from the Ribo-A system, and panels (e), (f), (g) and (h) so those from the Ribo-B system. Label “*c*_*i*_” is presented near the panel identifier. Two conformations are picked from either site *c*_2_ or *c*_3_. Red arrows indicate direction of the front view (see Figure S1 of SI).

Figures 2 and 3 demonstrates that sampling was done widely in the 3D RC space for both systems: ribocil from *c*_1_ was in the binding pocket, and ribocil from *c*_4_ was completely dissociated from the aptamer domain (Figures 3d and 3h). Ribocil from *c*_2_ was around the entrance of the binding pocket (Figures 3b and 3f), and ribocil from *c*_3_ was outside of the binding pocket with contacting the aptamer domain. Interestingly, we found ribocil on both the front and rear surfaces of the aptamer domain. This suggests that ribocil may enter the binding pocket from either surface of the tunnel.

The convergence check of the resultant distribution is argued in Section 5 of SI with a function *E*_*local*_(*i, j, k*), which was introduced originally in an earlier report (Higo et al., 2020a). This function is calculated at each 3D RC zone (*i, j, k*) in each iteration. Simply, when an inequality *E*_*local*_(*i, j, k*) < 0.25 is satisfied at a zone, we judged that *Q*_*cano*_ is determined accurately at the zone. Validity of this threshold value has been checked repeatedly (Higo, et al., 2020b; Higo et al., 2021; Hayami et al., 2021; Higo et al., 2022). Section 5 of SI demonstrated that *Q*_*cano*_(*i, j, k*) converged in both the bound and unbound regions of the 3D RC space in iterations 1–40 for both systems. We proceeded the simulation up to iteration 45 to increase the number of snapshots output.

### 3.2. Density of ribocil around aptamer domain and density tunnel

Figure 4 demonstrates the spatial density *ρ*_*CM*_ (***r***_*cube*_) of the ribocil’s centroid around the aptamer domain in the 3D real space, where the cube size was set to *L*_*c*_ *=* 2 Å (see Equation S1 of SI). The high-density region of *ρ*_*CM*_ *=* 0.0025 Å^−3^ (red contour region) was found only in the Ribo-B system. This region is referred to as “the highest-density spot” to state clearly that this density is the highest, and then we denote the highest density as *ρ*_*max*_ (i.e., *ρ*_*max*_ *=* 0.0025 Å^−3^). Figure 4c demonstrates that the highest density spot is in the binding pocket for the Ribo-B system. The region of *ρ*_*CM*_ *= ρ*_*max*_/10 (blue contour region) surrounded the highest density spot, and the regions of *ρ*_*CM*_ *= ρ*_*max*_/100 (magenta contour region) surrounded the blue contour region. This spatial pattern of *ρ*_*CM*_ relates to the funnel-like nature of Figure 2b. We note that the highest-density spot deviated by about 4 Å from the X-ray position (small black sphere) to a direction parallel to the red arrow in Figure 4d. We discuss this shift in the next subsection.

**Figure 4.**
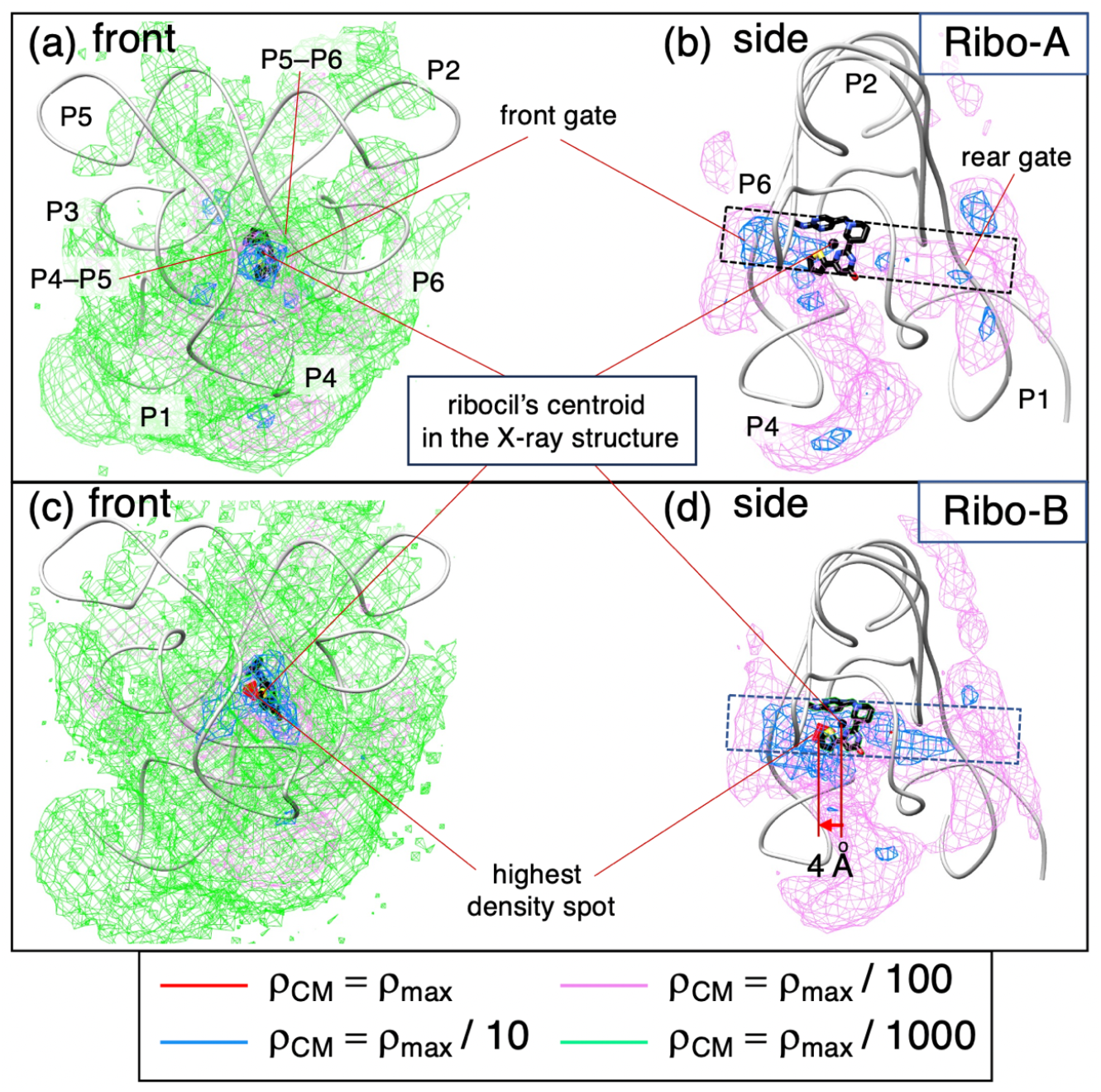
(a) Front and (b) side views of spatial density *ρ*_*CM*_(***r***_*cube*_) of ribocil A around the aptamer domain. Label P*i* (*i =* 1, …, 6) is an identifier assigned to P*i* helix. Label “P*i* − P*i* + 1” indicates junction region between P*i* and P*i* + 1. Similarly, front (c) and (d) side views of *ρ*_*CM*_(***r***_*cube*_) of the ribocil B. Iso-density levels are colored as indicated in inset, and *ρ*_*max*_ *=* 0.0025 Å^−3^. The density is normalized for both systems: 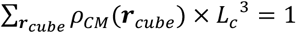. Light gray model represents aptamer domain, and green model does ribocil B in the X-ray structure (PDB ID: 5c45; P4 loop is modeled). Centroid of ribocil B in the X-ray structure is shown by black sphere. The structure of ribocil A is not determined experimentally. Black broken-line rectangle is density tunnel excavated from the front to rear surfaces of aptamer domain. Front and rear gates, from which ribocil enter the binding pocket, are shown in panels (a) and (b). See Figure S1 for P4–P6.

For the Ribo-A system, no spot of *ρ*_*CM*_ *= ρ*_*max*_ was found, and the binding pocket was partly occupied by the blue contours, although the volume for the blue contours was smaller than that for the Ribo-B system (Figure 4b). This feature of *ρ*_*CM*_ is more rugged than that for the Ribo-B system. Therefore, Figure 4 is consistent to Figure 2.

Remember that the experimental structure of the aptamer domain has a tunnel (Figure S1d and S1f) in either the apo or holo form. In fact, we found a density tunnel: The magenta contours (*ρ*_*CM*_ *= ρ*_*max*_/100) excavated the aptamer domain between the front and rear surfaces (the broken-line rectangle in Figures 4b and 4d). Therefore, ribocil can reach the binding position (the position in the X-ray complex) by passing gates on aptamer’s the front and rear surfaces. The gate (named “front gate”) on the front surface is a cleft formed by the P4–P5 junction and the P4–P5 junction of the aptamer domain (Figure 4a). The gate (“rear gate”) on the rear surface is a cleft opposite to the front gate (Figures 4b).

Because *ρ*_*CM*_ (***r***_*cube*_) in Figure 4 was presented with *L*_*c*_ *=* 2 Å, this figure does not show a fine structure of *ρ*_*CM*_ smaller than 2 Å. Then, to view the fine structure, we computed *ρ*_*CM*_ with *L*_*c*_ *=* 0.5 Å: The smaller the *L*_*c*_, the finer the spatial structure of *ρ*_*CM*_. Figure 5 indicates existence of a free-energy barrier (broken line) in both systems, whereas no clear free-energy barrier was seen in Figure 4. On the other hand, the number of snapshots detected in cubes decreases by decrement of *L*_*c*_. Although we examined *L*_*c*_ smaller than 0.5 Å, the spatial patterns of *ρ*_*CM*_ (***r***_*cube*_) became bumpy (data not shown).

**Figure 5.**
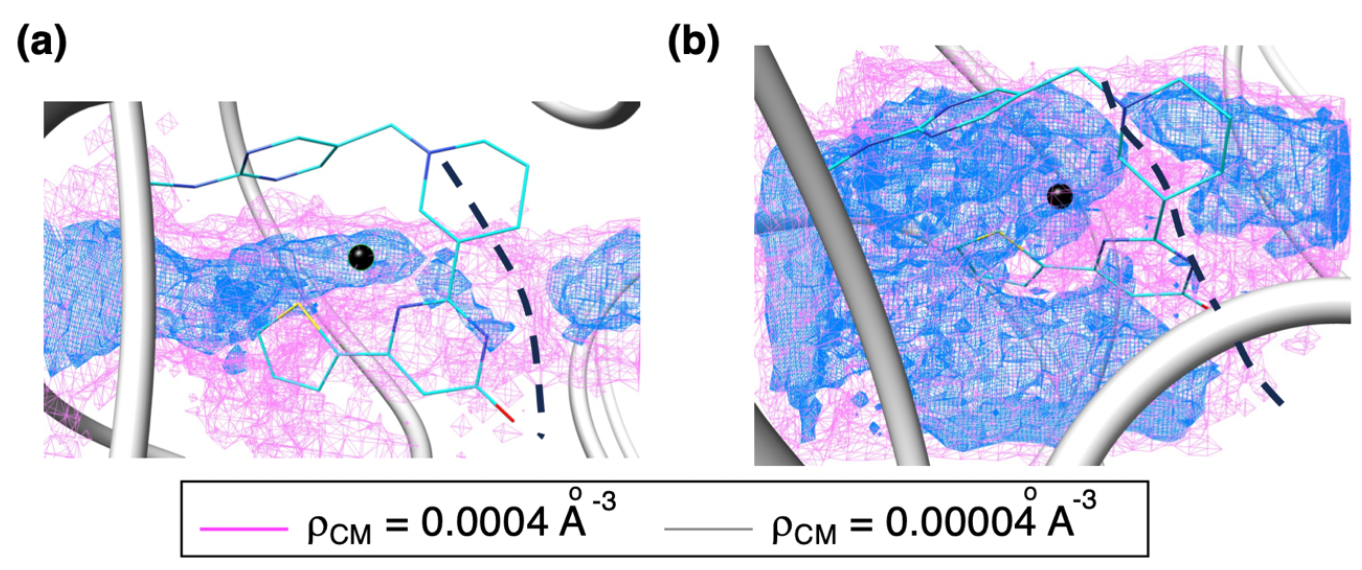
*ρ*_*CM*_ (***r***_*cube*_) with setting *L*_*c*_ *=* 0.5 Å focusing on the density tunnel (magenta contours of Figure 4) (a) Ribo-A system and (b) Ribo-B system.

### 3.3. Orientation of ribocil around aptamer domain

Figure 6 illustrates the spatial patterns of four quantities regarding the molecular orientation of ribocil B: < ***e***_↑_(***r***_*cube*_) >, < ***e***_←_(***r***_*cube*_) >, *SP*_↑_(***r***_*cube*_) and *SP*_←_(***r***_*cube*_). See Subsection 4.2 of SI for these quantities as well as 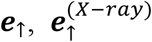 and 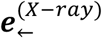. The patterns for ribocil A are presented in Figure S10 of SI. Now, we divide the density tunnel (broken-line rectangle in Figure 4) into two portions, front and rear portions of the density tunnel, as shown in Figure 6a.

**Figure 6.**
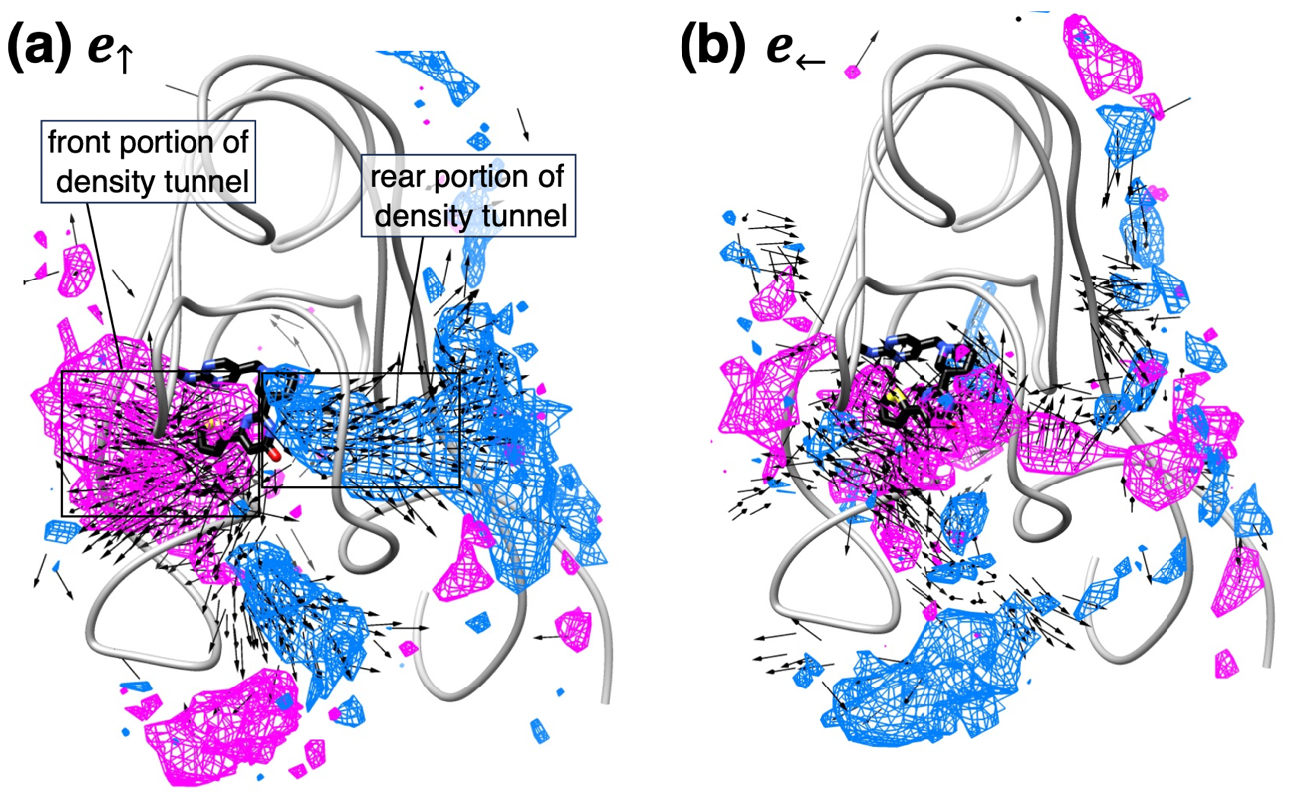
(a) < ***e***_↑_(***r***_*cube*_) > and *SP*_↑_(***r***_*cube*_) for Ribo-B system. (b) < ***e***_←_(***r***_*cube*_) > and *SP*_←_(***r***_*cube*_) for the Ribo-B system. In both panels, vectors < ***e***_*α*_(***r***_*cube*_) > (*α*=↑or ←) with | < ***e***_*α*_(***r***_*cube*_) > | ≥ 0.6 are shown by black arrows. Similarly, sites with *SP*_*α*_(***r***_*cube*_) ≥ 0.6 and *SP*_*α*_(***r***_*cube*_) ≤ −0.6 are shown, respectively, by magenta- and cyan-colored contours. Density tunnel is divided into the front and rear portions, which are indicated by broken-line rectangles. See also Figure S10 of SI for < ***e***_*α*_(***r***_*cube*_) > and *SP*_*α*_(***r***_*cube*_) of Ribo-A system.

Apparently, ***e***_↑_ in the front portion of the tunnel tended to be parallel to 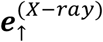 strongly as indicated by arrows of < ***e***_↑_(***r***_*cube*_) > and the magenta contours (*SP*_↑_(***r***_*cube*_) ≥ 0.6) of Figure 6a and Figure S10a. In contrast, ***e***_↑_ in the rear portion tended to be antiparallel to 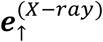 strongly as shown by arrows of < ***e***_↑_(***r***_*cube*_) > and the cyan contours (*SP*_↑_(***r***_*cube*_) ≤ −0.6). Interestingly, the magenta and blue contour regions switched from one to the other sharply at the boundary of the front and rear portions. Importantly, this boundary corresponds to the free-energy barrier of *ρ*_*CM*_ (***r***_*cube*_) (Figure 5). It is likely that the ligand rotation at the boundary accompanies a free-energy cost.

Next, we discuss the spatial patterns of ***e***_←_. Figure 6b and Figure S10b show that the regions of *SP*_←_(***r***_*cube*_) ≥ 0.6 dominated the density tunnel, whereas some small regions of *SP*_←_(***r***_*cube*_) ≤ −0.6 were found in the tunnel for both systems. Therefore, the spatial patterns for ***e***_←_did not show a clear transitional behavior at the boundary between the front and rear portions of the density tunnel. We concluded that the transitional motion of the ribocil’s molecular orientation at the boundary is related to flipping motions of ***e***_↑_as represented schematically in Figure S11.

Last of this subsection, we note that both vectors ***e***_↑_and ***e***_←_tended to be parallel to 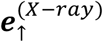 and 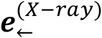, respectively, in the front portion of the density tunnel (magenta contours in both Figures 6 and S10). Therefore, we presume that there is a factor to maintain the molecular orientation. We discuss this point in the next subsection.

### 3.4. Stacking between aptamer domain and ribocil

Here we investigate a factor that stabilized complex conformations when ribocil was in the binding pocket. A study on the X-ray complex structures of the FMN riboswitch and some ligands reports that stacking between the ligand’s rings and the aptamer’s bases plays an important role to stabilize the complex structure (Rizvi et al., 2018). Interestingly, three aptamer’s bases A48, G62 and A85 were conserved in the stacking to bind to different ligands. Our simulation showed eight *π*-*π* stacking patterns (Table S4 of SI): Three of them were native stacking (Figures S7a–c of SI), and the other five were non-native stacking (Figures S7d–f). Those stacking patterns were found in both the Ribo-A and Ribo-B systems in our simulation when ribocil was in the binding pocket. Below, we analyzed the stacking quantitatively.

To pick up snapshots whose ribocil is in the binding pocket, we calculated root-mean-square-deviation of ribocil (*rmsd*_*rib*_) between a snapshot and the X-ray complex structure (PDB: 5c45). Remember that the aptamer part of snapshots had been superimposed to that of the X-ray structure (Subsection 4.1 of SI). We calculated *rmsd*_*rib*_ simply as: 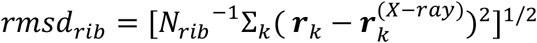, where ***r***_*k*_ is the position of the *k*-th heavy atom in ribocil in the snapshot, 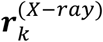 the corresponding atom in the X-ray complex structure, and *N*_*rib*_ the number of heavy atoms in ribocil. Collecting snapshots that satisfy an inequality *rmsd*_*rib*_ ≤ 5.0 Å, we assigned the stacking pairs to the collected snapshots using the method introduced in Subsection 4.3 of SI.

We computed the probability distribution function of the numbers of native and non-native stackings, 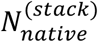 and 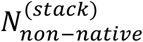, over the collected snapshots. Exact definition of 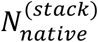 and 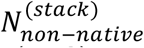 is given in Subsection 4.3 of SI. Figure 7a shows that the distribution 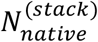 is similar between the two systems. In contrast, the non-native stacking was formed more in the Ribo-B system than in Ribo-A (Figure 7b). Therefore, Figures 7a and b suggest a possibility: The superior stability of ribocil B against ribocil A may be induced by the non-native stacking formed more in Ribo-B than in Ribo-A.

**Figure 7.**
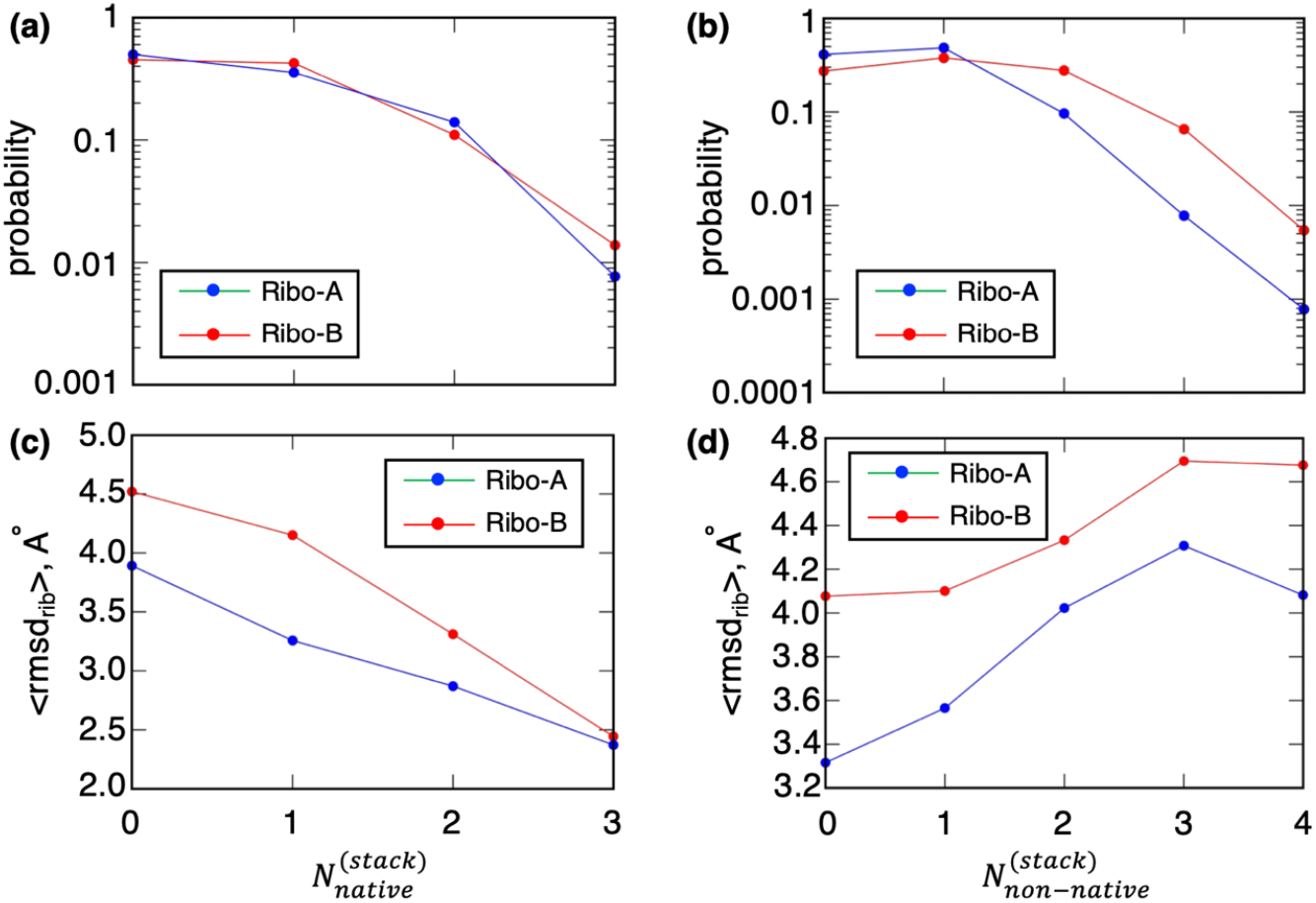
Distribution of 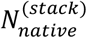 (a) and 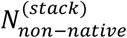 (b). See Subsection 4.3 of SI for definition of 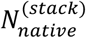 and 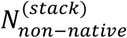. (c) < *rmsd*_*rib*_ > as a function of 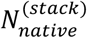 and (d) < *rmsd*_*rib*_ > as a function of 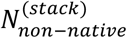. Snapshots with 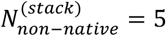 were not observed for both systems.

We investigate this possibility further. Remember that the highest-density spot of ribocil B deviated by about 4 Å from the X-ray position toward the entrance of the aptamer’s bonding pocket (see red arrow in Figure 4d). In fact, ribocil B deviated from the X-ray position toward the entrance (black arrows of Figure S8d–f of SI) when non-native stacking was formed. In contrast, ribocil B with the native stacking did not deviate largely from the X-ray position (Figures S8a–c). Thus, the deviation of the highest density spot from the X-ray position is likely to be caused by the formation of the non-native stacking. To make this visual impression quantitative, we calculated the average of *rmsd*_*rib*_ (i.e., < *rmsd*_*rib*_ >) as a function of 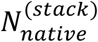 or 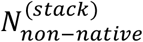, where the average was done using the statistical weight *w*_*i*_ assigned to each snapshot (see Subsection 4.1 of SI). Figure 7c demonstrates that the increment of 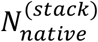 decreases < *rmsd*_*rib*_ >: The more the number of native stackings, the closer the conformation to the X-ray position. In contrast, Figure 7d shows that the increment of 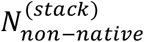 increased < *rmsd*_*rib*_ >. Therefore, we concluded that the snapshots contributing to the highest density spot are those stabilized by the non-native stacking.

This result presents a doubt that the superior stability of ribocil B against ribocil A may be derived artificially because the X-ray complex structure (PDB ID: 5c45) does not involve the non-native stacking patterns. However, *ρ*_*CM*_(***r***_*cube*_) in the vicinity of the X-tay position of ribocil B (Figure 5b) was larger than that for ribocil A. We note that the equilibrated probability of the system at a site is, in theory, determined simply by the potential energy at the site. Then, we calculated the probability assigned to a small volume around the X-ray position by:

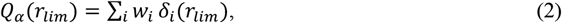

where *δ*_*i*_(*r*_*lim*_) = 1 when ***r***_*CM,i*_ is in a sphere (radius = *r*_*lim*_) centered at the ribocil’s centroid in the X-ray complex and *δ*_*i*_ (*r*_*lim*_) = 0 otherwise. The weight *w*_*i*_ is normalized: Σ_*i*_ *w*_*i*_ = 1, and *α* is an indicator of system: *α* = *A* for Ribo-A and *α* = *B* for Ribo-B.

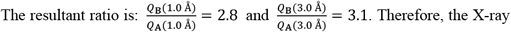

complex structure (PDB: 5c45) for the Ribo-B system is more stable than the complex structure for the Ribo-A system. Remember that the binding modes of ribocil A and B to the aptamer domain are almost identical (Howe, et al., 2016). We however, consider that this difference of stability between the two system is small. We discuss this point later.

### 3.5. Ribocil in the rear portion of the density tunnel

Ribocil in the rear portion of the density tunnel is not important biophysically because ribocil should overcome the free-energy barrier with varying the molecular orientation ***e***_↑_ largely, and then this overcoming motion makes the molecular binding slow. However, this result raises a question: Why does ***e***_↑_tend to be antiparallel to 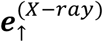 in the rear portion? If ***e***_↑_is parallel to 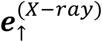, then ribocil can reach the X-ray position without the large orientational rearrangement, and consequently the free-energy barrier vanishes. In this subsection, we investigate ribocil’s conformations in the rear portion to the density tunnel.

We selected snapshots from the rear portion of the density tunnel with condition of *SP*_↑_ ≤ −0.9 and *SP*_←_ ≥ 0.5, and obtained 218 conformations. The first inequality specifies the snapshots to be almost antiparallel to 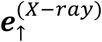. The second inequality imposes the snapshots so that the angle formed by ***e***_←_and 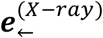 is 60° or smaller. We found that the majority (90 %) of them formed two hydrogen bonds between R3 of ribocil and G98 or A99 of the aptamer domain as well as between R1 and A102. We also found that intermolecular stacking was rarely formed (data not shown). Figure S12 of SI illustrates a typical snapshot with the hydrogen bonds. We conclude that snapshots with ***e***_↑_to be antiparallel to 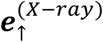 were stabilized by those hydrogen bonds. In other words, the free-energy barrier is a consequence of the hydrogen bonds when ribocil is in the rear portion of the density tunnel.

This inhibition mechanism for ribocil not to reach the X-ray position from the rear portion is interesting. We, however, could not argue only from the current simulation results if this mechanism is prepared evolutionally. We expect that the current simulation stimulates not only the computational field but also experimental or evolutionary biology.

## 4. Discussion

Encounter complexes were proposed in protein–ligand or protein–protein binding (Crowley et al., 2002; Xu et al., 2008; Kozakov, et al., 2014; Schilder & Ubbink, 2013). The present study also suggested that the ligand experiences encounter complexes before reaching the most favored site of the riboswitch (Figure 3). Starting from the unbound state, the ligand could contact almost the whole surface of riboswitch (green contours of Figure 4). Note that the green contour region has a lower free energy (potential of mean force; PMF) than the unbound region outside the green-contour region because PMF is defined by: *PMF* = −*RTln*[*ρ*_*CM*_ (***r***_*cube*_)], where *R* is gas constant. Then, the ligand found some stable sites (magenta contour sites) when moving on the surface. The density *ρ*_*CM*_(***r***_*cube*_) increased further in the vicinity of the front gate (blue contours in Figures 4b and d): The free energy decreased further around the front gate. Consequently, with proceeding to the binding pocket, the ligand was encapsulated.

We note this ligand orientational ordering is similar to the binding mechanism of a GPCR (human endothelin receptor type B) and a drug ligand (bosentan) reported in our earlier MD study (Higo et al., 2022). Namely, the ligand orientation ordered to enter the gate of the deep pocket of GPCR. The difference between two studies is: In the present study, the orientational ordering was brought by the ligand–receptor p-p stacking, whereas the ordering in the earlier study (Higo et al., 2022) was done by the interaction between the ligand and the long-disordered N-terminal tail of GPCR.

Both the apo and holo forms of the aptamer domain (PDB ID: 6wjr (Wilt et al., 2020) and 5c45 (Howe et al., 2015), respectively) have a tunnel, which excavates between the front and rear surfaces of the aptamer (Figure S2 of SI). In fact, the special density *ρ*_*CM*_ (***r***_*cube*_) showed that both ribocil A and B can pass through the tunnel (Figures 4b and 4d). Then, we referred to the tunnel as the density tunnel. The density tunnel was divided into the front and rear portions, and the binding pocket and the ligand binding sites were involved in the front portion (Figure 6a). Interestingly, a free-energy barrier existed at the boundary of the front and rear portions (Figure 5b). Further analysis showed that the molecular orientation < ***e***_↑_ > in the rear portion tended to be opposite to that in the front portion (Figure 6a) and 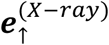. This means that the molecular orientation should turn completely at the boundary when the ligand passes the boundary from the rear to front portion (Figure S11 in SI). Thus, we presume that the main binding pathway is the approach from the front gate to the binding pocket.

It has been reported experimentally (Howe, et al., 2016) that the aptamer–ribocil B complex is considerably more stable than the aptamer–ribocil A complex is, whereas the binding forms of the two systems are almost identical. We showed that the free-energy landscape of the Ribo-B system is funnel like, and that for the Ribo-A system is rugged. These results agree qualitatively to the experimental report (Howe, et al., 2016). The highest density spot found for the Ribi-B system was located in the binding pocket (Figures 4c and 4d), and ***e***_↑_and ***e***_←_in the front portion of the density tunnel tend to be parallel to 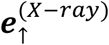 and 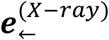 (Figure 6).

We, however, noticed from the detailed analysis that the highest density spot shifted by about 4 Å from the X-ray position toward the entrance of the binding pocket (red arrow in Figure 4d). One may consider that this shift is small and trivial, as found commonly in many simulations. However, the stacking analysis showed that the shift is not trivial. The shifted ribocil B (i.e., ribocil B at the highest density spot) was stabilized by the non-native stacking (Figure 7d). In fact, we showed that ribocil B in the vicinity of the X-ray position is supported by the native stacking (Figure 7c). Therefore, if the stability of the complex is conducted by the native stacking more than the non-native stacking, the highest density spots should emerge near the X-ray position.

We do not have a definite answer why the highest density spot shifted. However, two explanations are possible: One is the accuracy of the force field and the other is the efficiency of sampling. It has been reported that the currently used force field χ_OL3_ is an appropriate one to treat an RNA system by molecular simulation (Šponer et al., 2018). As we showed, the present computation provided good results except for the shift of the highest density spot, which relates to the non-native stacking. Thus, one possibility for this shift is inaccuracy of force field regarding the stacking. Because the *π*-*π* stacking interaction is determined quantum-mechanically (van Mourik & Hogan, 2016), errors may be introduced in the classical force field generation. Remember that ribocil B with the native stacking was well superimposed to that of the X-ray complex structure (Figures 8a–c and 9a–c): The native-like complex structure was sampled well in the simulation. This result suggests that the stability of the native-like complex structure increases if the stacking interaction is improved. The other possibility is insufficiency of conformational sampling. It is difficult to judge which the force-field inaccuracy or the sampling insufficiency is the determinant factor for the shift of the highest density spot. Because the thermodynamic stability of a many-body system is determined in a complicated manner, we cannot judge the determinant factor from the current simulation. We are planning to perform a simulation with adopting a new reaction-coordinate set.

Last, we discuss a positive role of the non-native stacking in the molecular binding. In general, the stacking is formed when two rings have appropriate positioning mutually, and it is likely that the non-native stacking is temporally formed when ribocil B is capsulated in the binding pocket, as we showed in this paper. Remember that U61 of the aptamer domain is located near the entrance of the binding pocket, and then ribocil B may be trapped shortly by U61 as shown in Figures S8d–f. These figures demonstrate that the non-native stacking between U61 and R4 of ribocil supports the molecular orientation ***e***_↑_to be parallel to 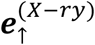. Then, riboil can reach the X-ray position by translational motions without rotation. In this scenario, the positioning of U61 is crucial to arrange the ribocil’s orientation to be advantageous for the molecular binding. Last, we emphasize that this scenario is less influenced by the accuracy of *π*-*π* stacking force field. Therefore, the present study is useful to understand the RNA-ligand binding scenario.

We uploaded important snapshots obtained from the current simulation to the Biological Structure Model Archive (BSM-Arc) (Bekker et al., 2020). The entry for this study is https://bsma.pdbj.org/entry/49.

## Acknowledgments

We are grateful to Dr. Kota Kasahara from Ritsumeikan University for implementation of the mD-VcMD algorithm to gromacs using the PLUMED plug-in. This work was supported by JSPS KAKENHI Grants No. 21K06052 (J.H.) and 20H03229 (N.K.), and performed in part under the Cooperative Research Program of the Institute for Protein Research, Osaka University, CR22-02 and CR-23-02. J.H. and K.N. acknowledge support by the HPCI System Research Project (Project IDs: hp220002, hp220015, hp220022, hp230003, and hp230011). J.H., I.F., N.K., and Y.F. are supported by the Project Focused on Developing Key Technology for Discovering and Manufacturing Drugs for Next-Generation Treatment and Diagnosis (2018–2021 and 2021–) from AMED and the Japan Biological Informatics Consortium (JBiC). GA-mD-VcMD was performed on TSUBAME3.0 supercomputers at the Tokyo Institute of Technology. We used UCSF Chimera ver. 1.15 for drawing molecular structures and gnuplot for graphs. Other analyses were done using homemade programs.

## Authors’ contributions

J.H., N.K., I.F., and Y.F. designed the research plan; J.H. developed the mD-VcMD algorithm; G.J.B. and N.K. generated the molecular system for simulation and input files for gromacs, and then performed preparatory simulations; J.H. performed the main part of the simulation and analyses; I.F. and Y.F. performed some important parts of analyses; J.H. wrote the paper.

## Data availability

The datasets used or analyzed during the current study are available from the corresponding author on reasonable request. Several important snapshots during the current simulation were available in the BSM-Arc (https://bsma.pdbj.org/entry/49).

## Supplementary Information (SI)

**Figure S1:**
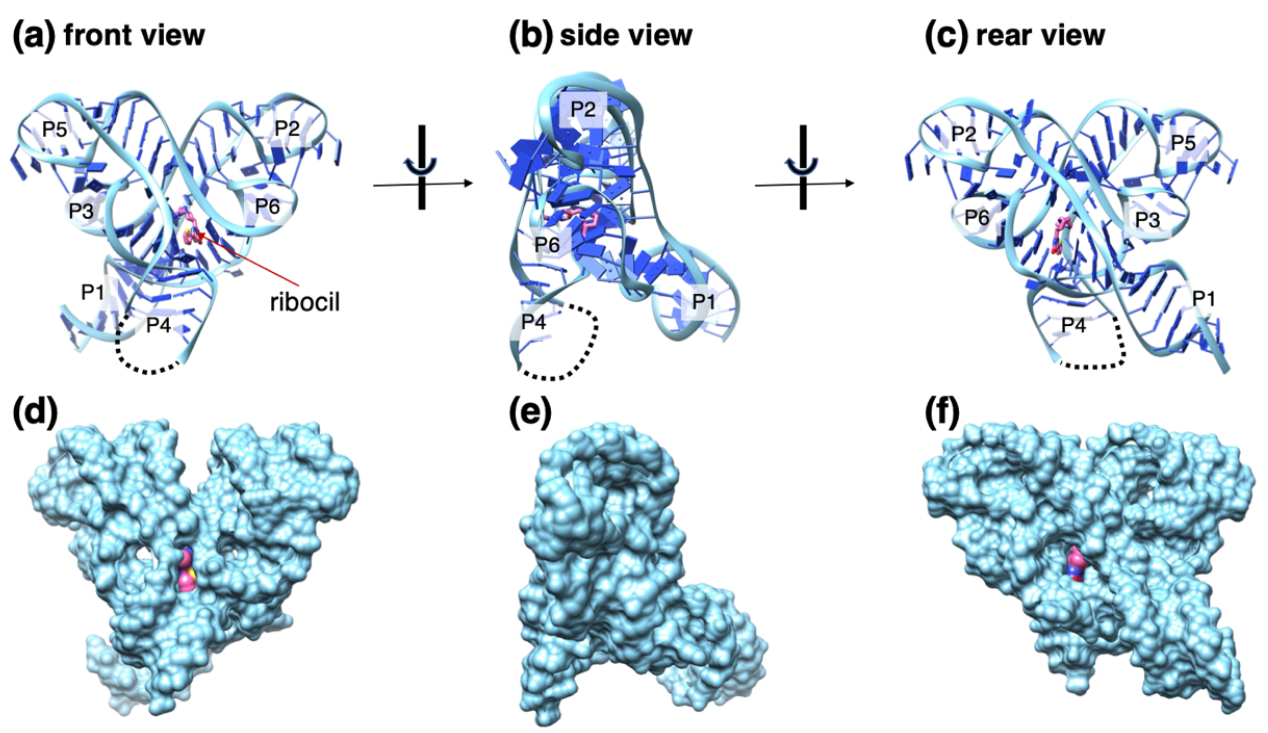
(a) Front, (b) side and (c) rear views of complex structure (PDB: 5c45) of the FMN riboswitch aptamer domain (cyan) and ribocil B (magenta) (Howe et al., 2016). Panels (d), (e), and (f) are, respectively, the front, side, and rear views with space filling model. Label P*i* (*i* = 1, …, 6) in panels (a)–(c) is an identifier assigned to P*i* helix (Wilt et al., 2020; Howe et al., 2016). Part of P4 is undetermined in the PDB structure, which is presented by broken line. Panels (d) and (f) show that the aptamer domain has a deep pocket, to which ribocil B binds.

**Figure S2:**
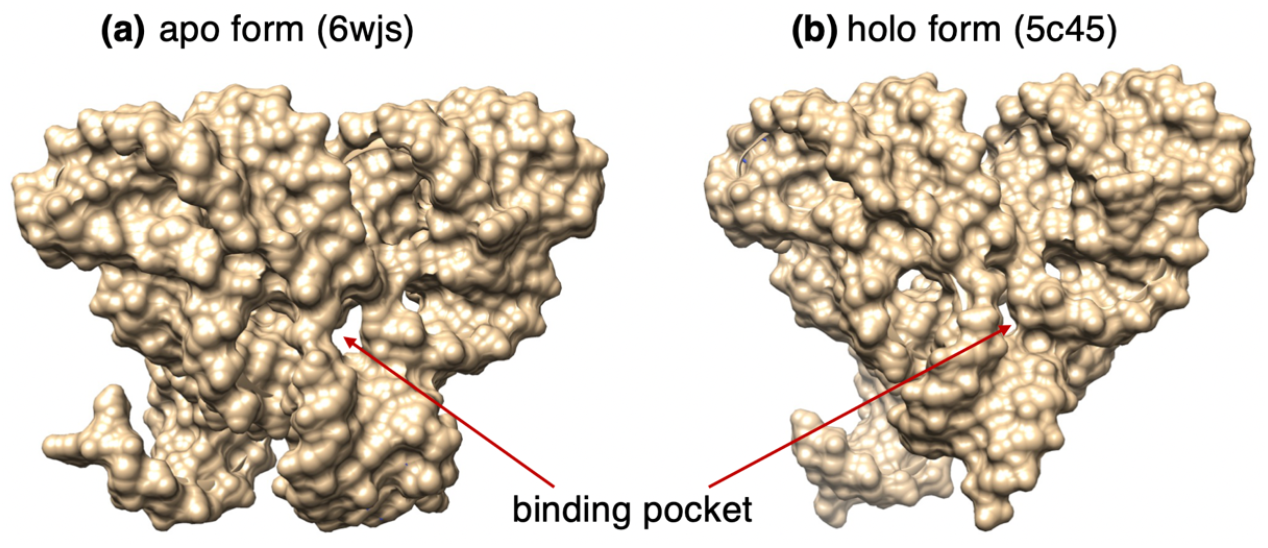
Aptamer domain (space filling model) of the FMN riboswitch of (a) apo form (PDB ID: 6wjr) and (b) holo form (PDB ID: 5c45). Both panels are illustrated from the front direction (see Figure S1) after removing ions and ligand. The binding pocket is indicated by arrows, which is filled by ligand in the apo form.

### Section 1: Simulation system and initial conformation

Here we explain the procedure to generate the two simulation systems: Ribo-A (ligand = ribocil A) and Ribo-B (ribocil B). The structure of the aptamer domain was taken from an X-ray crystallography (PDB entry = 6wjr; 2.7 Å resolution), which is an apo form of the aptamer domain of the FMN riboswitch (Wilt et al., 2020). There are other apo forms for the aptamer domain in PDB: 2yif (3.298 Å) (Vicens et al., 2011) and 6wjs (2.8 Å) (Wilt et al., 2020). Then, we selected the X-ray structure with the smallest resolution (i.e., 6wjr) of the three.

The ligands, ribocil A and B, are optical isomers mutually (Figures 1a and b) and flexible because the dihedral bonds of ligands are rotatable. The structures and the force field parameters for the ligands are modeled by us as explained in the main text.

We put the aptamer domain at the center of a periodic boundary box filled by water molecules, whose size was 97.9582 Å × 97.9582 Å × 97.9582 Å. To generate an unbound conformation for the initial conformation of simulation, ribocil A (or B) was put at the corner of the periodic box as shown in Figure S3a of SI. In immersing the aptamer domain and ribocil in the box, water molecules overlapping to the aptamer domain and ribocil were removed. To determine the number of Mg ions to be introduced in the system, we checked 19 X-ray structures regarding the aptamer domain (entries = 2yie, 2yif, 3f2q, 3f2t, 3f2w, 3f2x, 3f2y, 3f4e, 3f4g, 3f4h, 3f30, 5c45, 5kx9, 6bfb, 6dn1, 6dn2, 6dn3, 6wjr, and 6wjs) available in 2021. Then, the number of the maximum Mg ions in the X-ray structures was 15 from the entry of 3f4g, which is a complex structure of the aptamer domain and riboflavin. This may mean that the aptamer domain has 15 sites bindable to Mg ions. Thus, we introduced 15 Mg ions at the 15 Mg sites found in 3f4g. K and Cl ions were also introduced to set the ionic concentration to a physiological level (0.1 M) with condition that the net charge of the entire system should be neutralized. Water molecules overlapping to the introduced ions were removed. Consequently, the number of atoms involved in the systems was 92,628: 3,572 atoms for the aptamer, 49 ligand atoms, 29,602 water molecules, 15 Mg ions, 133 K ions and 53 Cl ions. The number of constituent atoms was the same between the two systems because ribocil A and B are isomers and the number of water molecules to be removed was the same when introducing ribocil.

The above conformations were energy-minimized followed by an NVT simulation (*T* = 300 *K*). Then, an NPT simulation (*T* = 300 *K* and *P* = 1 *atm*) was performed to relax the system. Figure S3b of SI illustrates the resultant conformation from the NPT simulation, and this conformation was used for the initial conformation of the current mD-VcMD simulation. The resultant box size was 97.3511 Å × 97.3511 Å × 97.3511 Å and 97.3304 Å × 97.3304 Å × 97.3304 Å for the Ribo-A and Ribo-B systems, respectively. Ribocil was still near the corner of the periodic box. The force-field parameters used for the simulations were explained in the main text.

**Figure S3:**
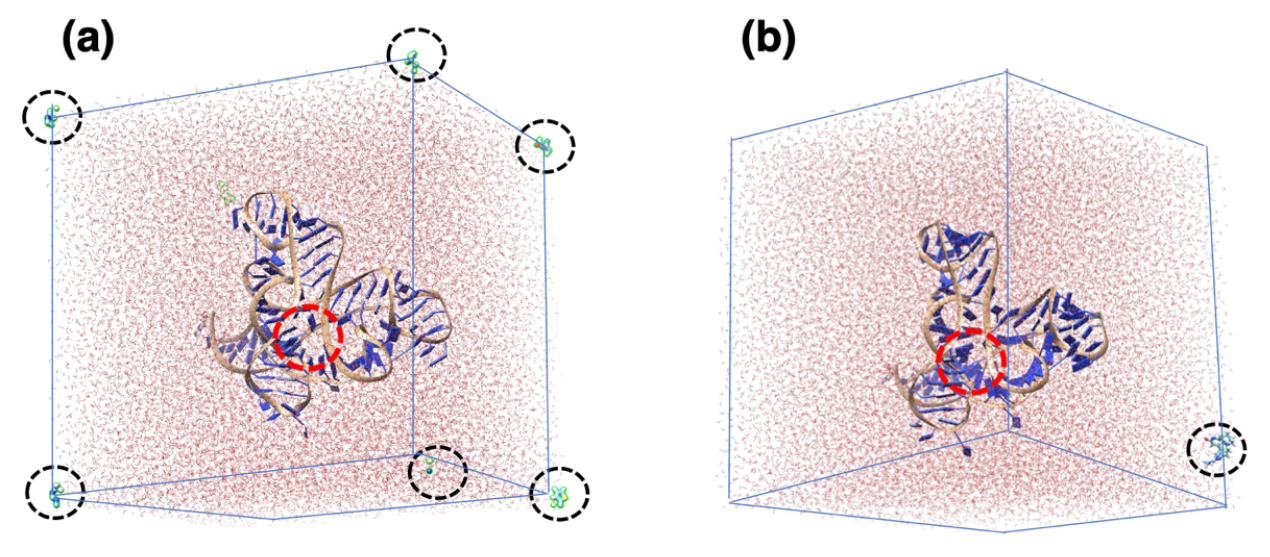
(a) Ribo-A system. The aptamer domain is put at the center of the periodic box, and ribocil A is put at the corner of the box, which is the reason why ribocil A is split into pieces (black broken-line circles). Red circle represents the ligand-binding site of the aptamer domain. Ribo-B system is not shown because it is very close to Ribo-A system. Note that the split segments of ribocil A are treated as a single molecule in simulation under the periodic boundary condition. (b) Conformation of Ribo-A system after NPT simulation. Black broken-line circle indicates ribocil A. Although the Ribo-B system is not shown, ribocil B is also near a corner of the box.

### Section 2: Definition of a reaction coordinate

Definition of a single reaction coordinate (RC), denoted as *λ*^(*h*)^, is explained here. Consider two atom groups 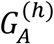 and 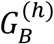 in a molecular system (Figure S4 of SI). Then, *λ*^(*h*)^ is defined by the distance between the centroids of 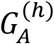 and 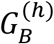. The actual set of 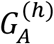 and 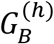 for the present study is explained later. Although we used centroids of atom groups to define RCs in this paper, RCs can be arbitrary quantities, which are expressed by the system’s coordinates in theory.

**Figure S4:**
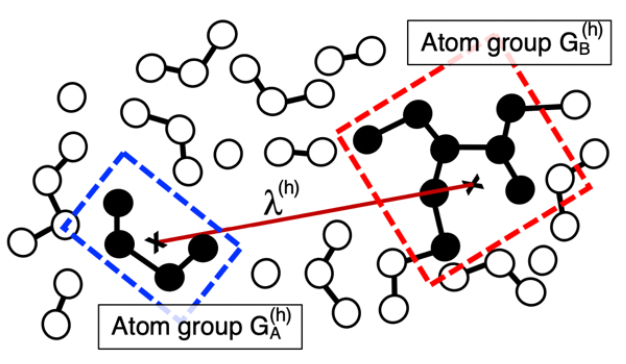
To define an RC, *λ*^(*h*)^, two atom groups 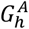 and 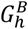 are introduced, which are shown by broken-line rectangles in blue and red, respectively, whose constituent atoms are shown by black filled circles. Centroid of each atom group is presented by a cross. *λ*^(*h*)^ is the distance (brown line) between the two centroids.

### Section 3. Atom groups to define reaction coordinates

The method to define an RC is given in Section 2 of SI. Here, we introduce three RCs: *λ*^(*h*)^ (*h* = *α, β, γ*), which are used for the present study. Figure 1c of the main text visualize the atom groups 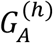 and 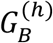 to identify the three RCs. Table S1 lists the contents of the atom groups.

**Table S1.** Atom groups to define three RCs ^a)^

| RC | atom group | contents |
| --- | --- | --- |
| $\lambda^{(\alpha)}$ | $G_A^{(\alpha)}$ | U34, G35, C44, A45, A87, G88, C95, U96 of aptamer domain |
| | $G_B^{(\alpha)}$ | Segment of ribocil as indicated in Figure 1c |
| $\lambda^{(\beta)}$ | $G_A^{(\beta)}$ | A48, A49 of aptamer domain |
| | $G_B^{(\beta)}$ | Segment of ribocil as indicated in Figure 1c |
| $\lambda^{(\gamma)}$ | $G_A^{(\gamma)}$ | U61, G62 of aptamer domain |
| | $G_B^{(\gamma)}$ | G84, A85 of aptamer domain |

<sup>a)</sup> When an atom group is assigned to a part of the aptamer domain (i.e., $G_A^{(\alpha)}$ , $G_A^{(\beta)}$ , $G_A^{(\gamma)}$ , and $G_B^{(\gamma)}$ ), the atom group involves only main-chain atoms. Figure 1c of main text visualizes atom groups and three RCs.

**Table S2.**
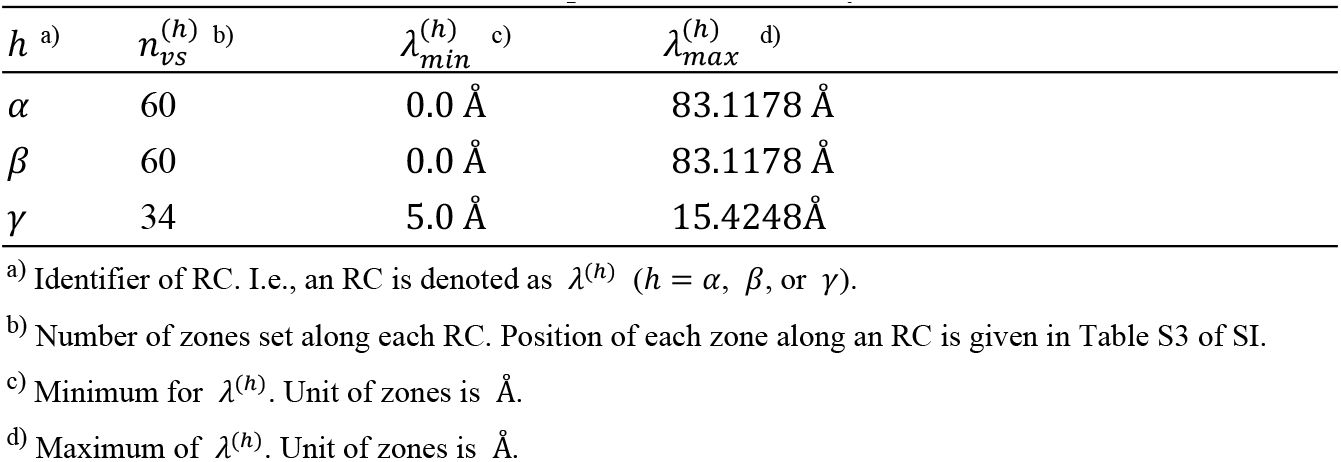
Parameters for 3D-RC space constructed by *λ*^(*α*)^, *λ*^(*β*)^ and *λ*^(*γ*)^

**Table S3.**
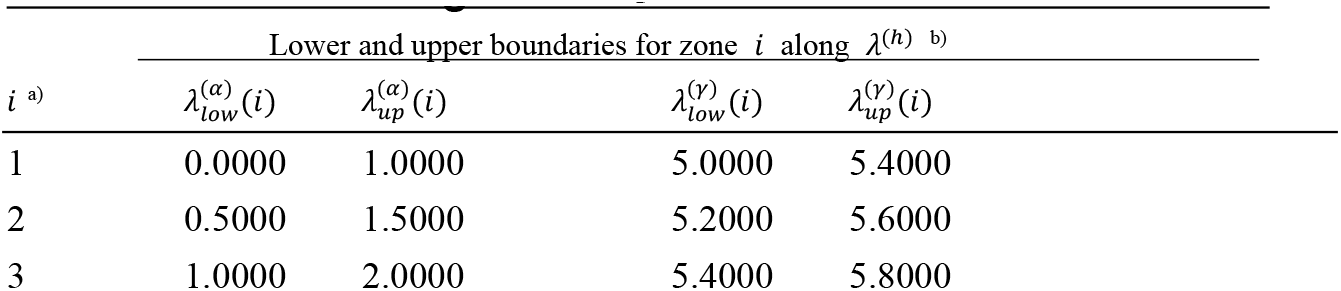
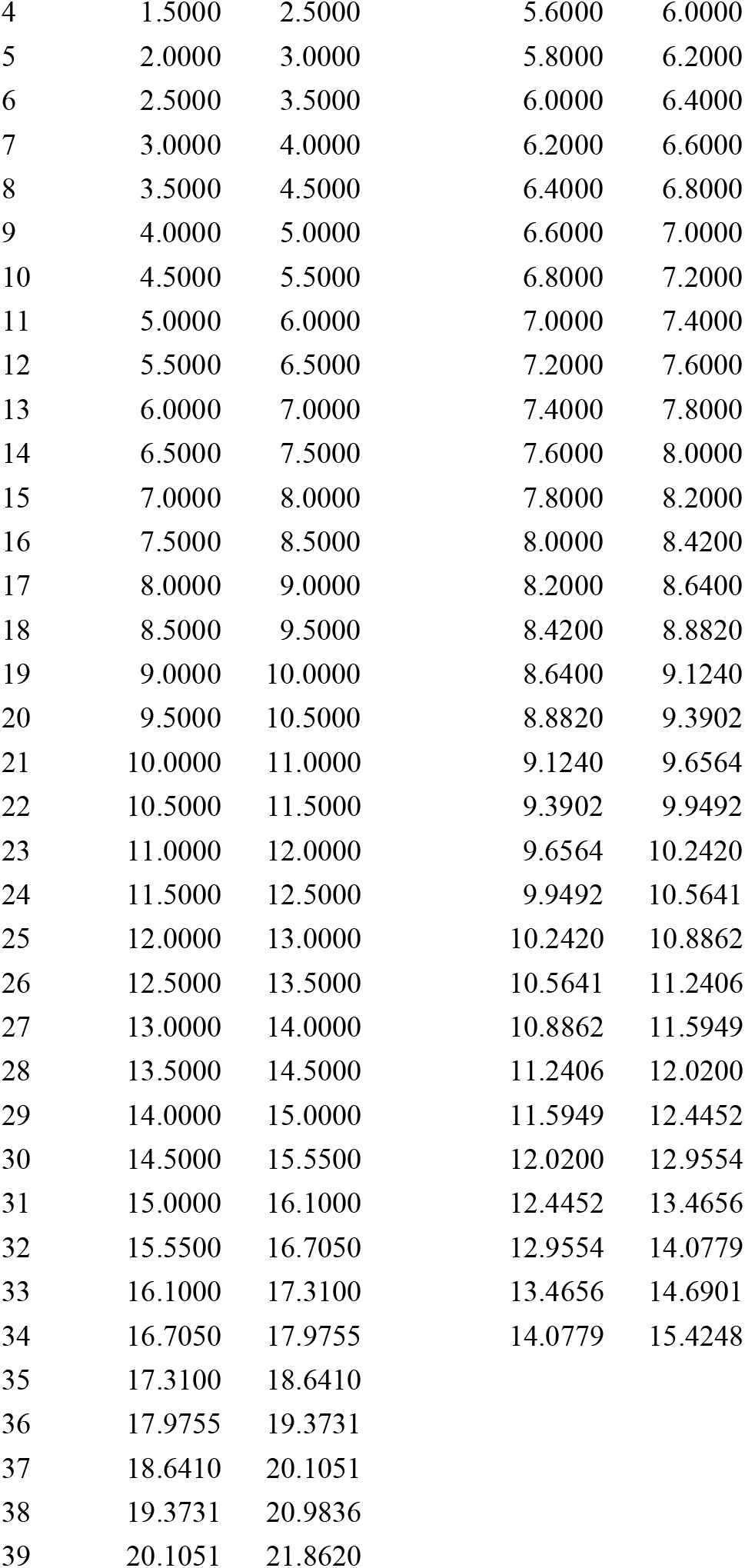
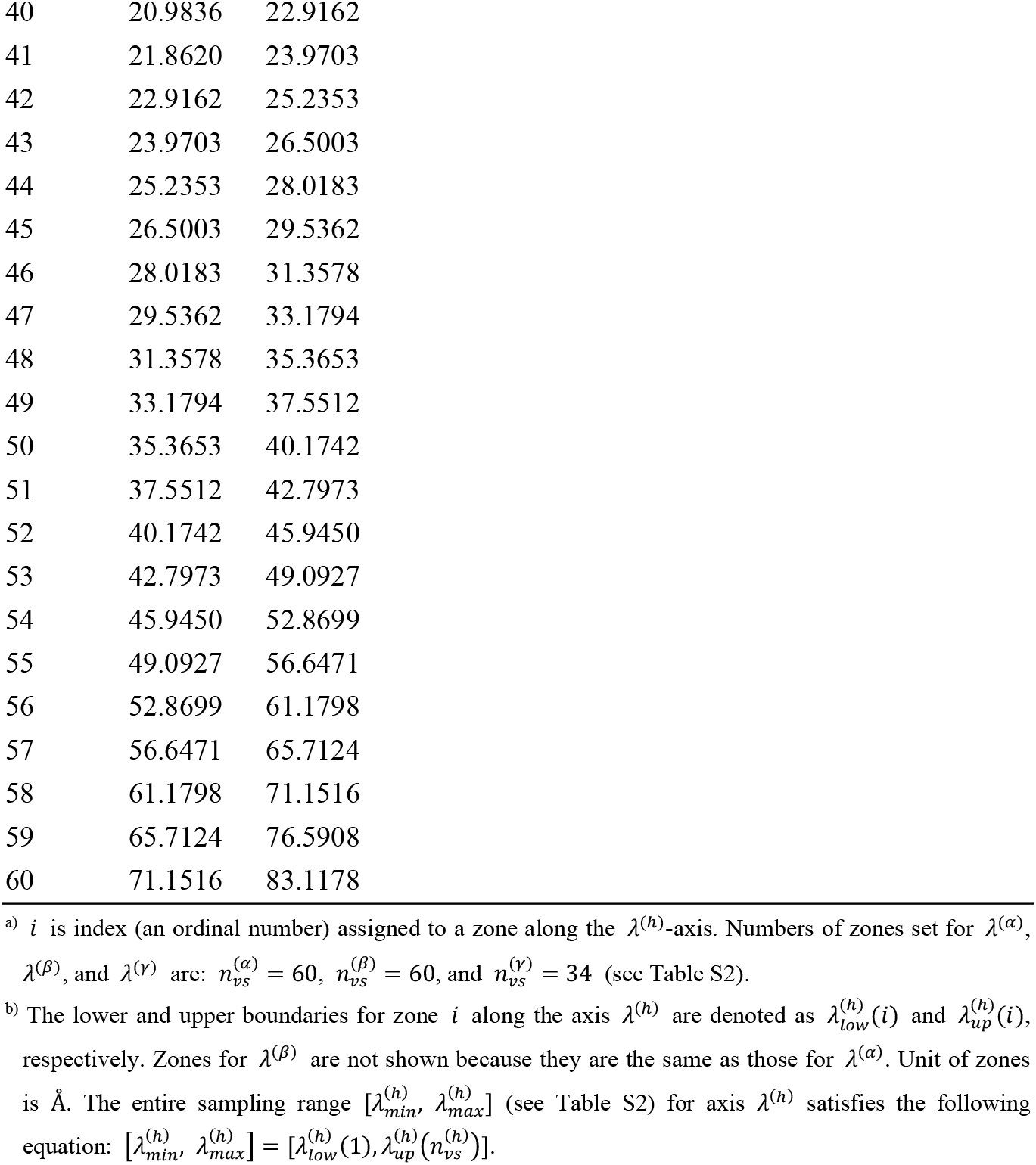
Zones along each RC.

| $i$ <sup>a)</sup> | Lower and upper boundaries for zone $i$ along $\lambda^{(h)}$ <sup>b)</sup> | | | |
| --- | --- | --- | --- | --- |
| | $\lambda_{low}^{(\alpha)}(i)$ | $\lambda_{up}^{(\alpha)}(i)$ | $\lambda_{low}^{(\gamma)}(i)$ | $\lambda_{up}^{(\gamma)}(i)$ |
| 1 | 0.0000 | 1.0000 | 5.0000 | 5.4000 |
| 2 | 0.5000 | 1.5000 | 5.2000 | 5.6000 |
| 3 | 1.0000 | 2.0000 | 5.4000 | 5.8000 |
| 4 | 1.5000 | 2.5000 | 5.6000 | 6.0000 |
| 5 | 2.0000 | 3.0000 | 5.8000 | 6.2000 |
| 6 | 2.5000 | 3.5000 | 6.0000 | 6.4000 |
| 7 | 3.0000 | 4.0000 | 6.2000 | 6.6000 |
| 8 | 3.5000 | 4.5000 | 6.4000 | 6.8000 |
| 9 | 4.0000 | 5.0000 | 6.6000 | 7.0000 |
| 10 | 4.5000 | 5.5000 | 6.8000 | 7.2000 |
| 11 | 5.0000 | 6.0000 | 7.0000 | 7.4000 |
| 12 | 5.5000 | 6.5000 | 7.2000 | 7.6000 |
| 13 | 6.0000 | 7.0000 | 7.4000 | 7.8000 |
| 14 | 6.5000 | 7.5000 | 7.6000 | 8.0000 |
| 15 | 7.0000 | 8.0000 | 7.8000 | 8.2000 |
| 16 | 7.5000 | 8.5000 | 8.0000 | 8.4200 |
| 17 | 8.0000 | 9.0000 | 8.2000 | 8.6400 |
| 18 | 8.5000 | 9.5000 | 8.4200 | 8.8820 |
| 19 | 9.0000 | 10.0000 | 8.6400 | 9.1240 |
| 20 | 9.5000 | 10.5000 | 8.8820 | 9.3902 |
| 21 | 10.0000 | 11.0000 | 9.1240 | 9.6564 |
| 22 | 10.5000 | 11.5000 | 9.3902 | 9.9492 |
| 23 | 11.0000 | 12.0000 | 9.6564 | 10.2420 |
| 24 | 11.5000 | 12.5000 | 9.9492 | 10.5641 |
| 25 | 12.0000 | 13.0000 | 10.2420 | 10.8862 |
| 26 | 12.5000 | 13.5000 | 10.5641 | 11.2406 |
| 27 | 13.0000 | 14.0000 | 10.8862 | 11.5949 |
| 28 | 13.5000 | 14.5000 | 11.2406 | 12.0200 |
| 29 | 14.0000 | 15.0000 | 11.5949 | 12.4452 |
| 30 | 14.5000 | 15.5500 | 12.0200 | 12.9554 |
| 31 | 15.0000 | 16.1000 | 12.4452 | 13.4656 |
| 32 | 15.5500 | 16.7050 | 12.9554 | 14.0779 |
| 33 | 16.1000 | 17.3100 | 13.4656 | 14.6901 |
| 34 | 16.7050 | 17.9755 | 14.0779 | 15.4248 |
| 35 | 17.3100 | 18.6410 |  |  |
| 36 | 17.9755 | 19.3731 |  |  |
| 37 | 18.6410 | 20.1051 |  |  |
| 38 | 19.3731 | 20.9836 |  |  |
| 39 | 20.1051 | 21.8620 |  |  |

|  |  |  |
| --- | --- | --- |
| 40 | 20.9836 | 22.9162 |
| 41 | 21.8620 | 23.9703 |
| 42 | 22.9162 | 25.2353 |
| 43 | 23.9703 | 26.5003 |
| 44 | 25.2353 | 28.0183 |
| 45 | 26.5003 | 29.5362 |
| 46 | 28.0183 | 31.3578 |
| 47 | 29.5362 | 33.1794 |
| 48 | 31.3578 | 35.3653 |
| 49 | 33.1794 | 37.5512 |
| 50 | 35.3653 | 40.1742 |
| 51 | 37.5512 | 42.7973 |
| 52 | 40.1742 | 45.9450 |
| 53 | 42.7973 | 49.0927 |
| 54 | 45.9450 | 52.8699 |
| 55 | 49.0927 | 56.6471 |
| 56 | 52.8699 | 61.1798 |
| 57 | 56.6471 | 65.7124 |
| 58 | 61.1798 | 71.1516 |
| 59 | 65.7124 | 76.5908 |
| 60 | 71.1516 | 83.1178 |

<sup>a)</sup> $i$ is index (an ordinal number) assigned to a zone along the $\lambda^{(h)}$ -axis. Numbers of zones set for $\lambda^{(\alpha)}$ , $\lambda^{(\beta)}$ , and $\lambda^{(\gamma)}$ are: $n_{vs}^{(\alpha)} = 60$ , $n_{vs}^{(\beta)} = 60$ , and $n_{vs}^{(\gamma)} = 34$ (see Table S2).
<sup>b)</sup> The lower and upper boundaries for zone $i$ along the axis $\lambda^{(h)}$ are denoted as $\lambda_{low}^{(h)}(i)$ and $\lambda_{up}^{(h)}(i)$ , respectively. Zones for $\lambda^{(\beta)}$ are not shown because they are the same as those for $\lambda^{(\alpha)}$ . Unit of zones is Å. The entire sampling range $[\lambda_{min}^{(h)}, \lambda_{max}^{(h)}]$ (see Table S2) for axis $\lambda^{(h)}$ satisfies the following equation: $[\lambda_{min}^{(h)}, \lambda_{max}^{(h)}] = [\lambda_{low}^{(h)}(1), \lambda_{up}^{(h)}(n_{vs}^{(h)})]$ .

### Section 4. Analyzed quantities

#### Subsection 4.1. Structural superposition and spatial density of ligand

The simulation produced 460,800 snapshots for each of the Ribo-A and Ribo-B systems as mentioned in the main text. The mD-VcMD assigns an equilibrated thermodynamic weight (statistical weight) at a simulation temperature (300 K in the present study) to all the snapshots (Higo et al., 2020a). Here, we denote the weight assigned to snapshot *i* as *w*_*i*_.

In this section, we introduce some physical quantities to analyze the ligand–receptor binding process. However, before introducing these quantities, we explain a structural superposition procedure to present a framework useful for the analysis. In an MD simulation, the molecules move translationally and rotationally in the periodic boundary box. Therefore, after the simulation, to analyze the behavior of the ligand around the receptor, the translational and rotational motions should be eliminated from snapshots. For this purpose, the aptamer domain of each snapshot was superimposed (mass-weighted superposition) on that of the initial conformation of simulation (Figure S3b). The other portions than the aptamer domain of the system (i.e., ribocil, water molecules and ions) were moved according to the superposition. This superposition corresponds to definition of a unified coordinate system (a body-fixed coordinate system) on the aptamer domain. This superposition was applied to all the snapshots. Note that any physical quantity reported in this paper is computed using the superimposed coordinates, although we do not mention it explicitly.

Next, the 3D real space (not the 3D-RC space) was divided into small cubes whose volume is *L*_*c*_ × *L*_*c*_ × *L*_*c*_, where *L*_*c*_ is the length of sides of the cube (not a zone given in Table S3). The position of a cube is represented by its cube–center ***r***_*cube*_ in the 3D real space. The position of the centroid (center of mass) of the ribocil A or B was referred to as ***r***_*CM*_. When we specify that ***r***_*CM*_ is computed from snapshot *i*, then we denote it as ***r***_*CM,i*_. Assigning the ligand’s centroid of all snapshots to cubes, we calculated the spatial density *ρ*_*CM*_(***r***_*cube*_) of the ligand’s centroid in the 3D real space as:

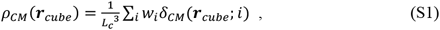

where *δ*_*CM*_ (***r***_*cube*_; *i*) is a delta function defined as:

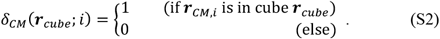

Last, we normalized *ρ*_*CM*_ (***r***_*cube*_) as:

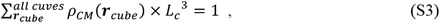

where the summation is taken over all cubes for each system.

#### Subsection 4.2. Ligand orientation around receptor

We analyze ribocil’s molecular orientation around the aptamer domain by the following procedure. First, for convenience, the four rings in ribocil are named as R1–R4 as shown in Figure S5. We defined the ligand’s orientation by two vectors. One is a unit vector ***e***_←_ parallel to a vector ***v***_←_, which is defined from the centroid of R3 to that of R2: ***e***_←_ = ***v***_←_/|***v***_←_|. The other is a unit vector ***e***_↑_ parallel to a vector ***v***_↑_, which is defined from the centroid of R2 and R3 to that of R1 and R4: ***e***_↑_ = ***v***_↑_/|***v***_↑_|. The reason why we introduced two vectors is: a single vector cannot specify the ligand orientation because the ligand can rotate around the single vector. Figure S5 presents ***e***_←_ and ***e***_↑_ visually.

In general, to assign a quantity to a site in the 3D real space, we use the cube introduced in the above subsection. Referring the quantity computed from snapshot *i* as to Ω_*i*_, the field quantity regarding Ω is defined as:

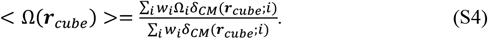

< Ω(***r***_*cube*_) > is the expectation value of Ω in cube at ***r***_*cube*_, where Ω can be either a scalar or vector. By setting as Ω = ***e***_←_ or ***e***_↑_, the orientation field < ***e***_←_(***r***_*cube*_) > or < ***e***_↑_(***r***_*cube*_) > is computed.

The quantities < ***e***_←_(***r***_*cube*_) > and < ***e***_↑_(***r***_*cube*_) > indicate how the molecular orientation is ordered in each cube. However, these quantities do not teach how the molecular orientation of ribocil has a similarity with that of ribocil in the native complex (the X-ray complex structure). Then, to investigate this orientational similarity, we introduced the following scalar product:

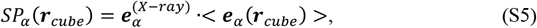

where *α* =← or ↑, and 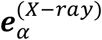 is the molecular orientation vector of ribocil in the X-ray complex (PDB: 5c45). The larger the *SP*_*α*_(***r***_*cube*_) in a cube at ***r***_*cube*_, the more similar the < ***e***_*α*_ (***r***_*cube*_) > to 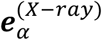 in a range of −1 ≤ *SP*_*α*_ (***r***_*cube*_) ≤ +1.

**Figure S5:**
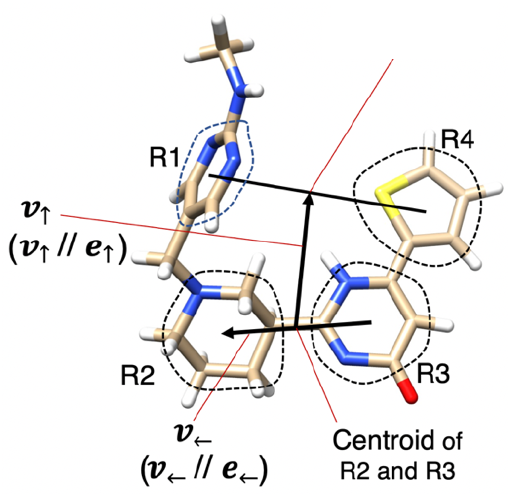
Ligand’s molecular orientation vectors ***e***_←_ and ***e***_↑_, whose exact definition is explained in text of SI. Four rings of ribocil, indicated by broken-line circles, are named as R1–R4. The centroid of a ring is calculated with eliminating hydrogen atoms, and the oxygen atom of Ring 3 is also eliminated.

#### Subsection 4.3. Ligand–receptor *π*-*π* stacking

The complex of ribocil B and the aptamer domain (PDB: 5c45) shows three intermolecular *π*-*π* stacking patterns (face-to-face stacking), which are referred to as “native stacking” in this paper. Figure S6a of SI demonstrates these stacking patterns: A48–R4, G62–R1 and A85–R3. We also presented two bases A49 and U61 in blue because they participated in ring–stacking (non-native stacking) other than the native stacking as explained in the *Results and Discussion* of the main text.

**Figure S6:**
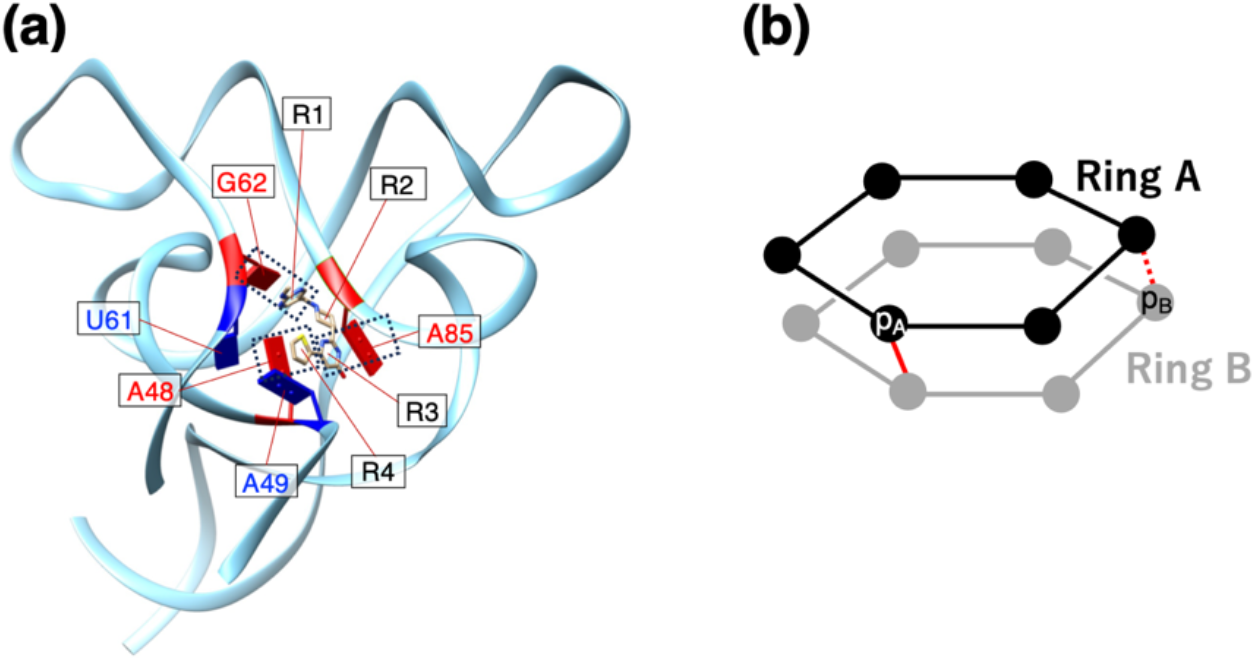
(a) Complex structure (PDB: 5c45). Three native ring-stacking patterns are indicated by broken-line rectangles, where A48, G62, and A85 are colored in red. Two bases colored in blue (A49 and U61) are those which participated to non-native stacking in simulation. Detailed discussion for the non-native stacking is presented in main text. L4 region of the aptamer domain is not shown in this panel because L4 is not determined in the X-ray complex structure. (b) Scheme of *π*-*π* stacking: Black and gray spheres are atoms belonging to Ring A and Ring B, respectively. Atoms *p*_*A*_ and *p*_*B*_ as well as red solid and red broken lines are mentioned in text of SI.

Now, we explain the procedure to detect intermolecular stacking in a snapshot. In principle, stacking is induced quantum chemically (van Mourik et al., 2016). On the other hand, a force field used in molecular dynamics is designed to mimic classical mechanically the stacking geometry. Here, we defined the stacking by relative positioning of two rings. Figure S6b displays schematically two rings, which are named Ring A (black) and Ring B (gray) for convenience. We consider only the heavy atoms in the ring backbone ignoring other atoms. Then, we calculated the minimum distance from an atom of Ring A (the atom marked “*p*_*A*_” for instance) to all backbone heavy atoms in Ring B in Figure S6b. Assume that the red solid line corresponds to the minimum distance 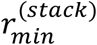. If the minimum distance was smaller than 3.6 Å, we judged that the atom “*p*_*A*_” is contacting to Ring B. We repeated this calculation for all backbone heavy atoms in Ring A and denote the number of atomic contacts from Ring A to Ring B as 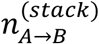. The threshold of 3.6 Å was set by viewing the native stacking in the native complex (PDB: 5c45): 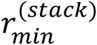 for the native stacking was about 3.5 Å. Then we set the threshold by adding a tolerance of 0.1 Å: 3.5 Å + 0.1 Å.

Similarly, we calculated the number of contacts from Ring B to Ring A: The minimum distance from atom marked “*p*_*B*_” in Ring B to the backbone-heavy atoms of Ring A was calculated and checked if the minimum distance is smaller than 3.6 Å. With repeating this procedure for all backbone heavy atoms in Ring B, we calculated the number of contacts, 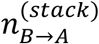, from Ring B to Ring A.

Last, we judged that Ring A and Ring B are stacking mutually if 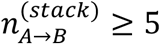 and 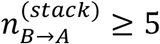. The rings in Figure S6b are six-membered rings. In the current system, ribocil involves five- and six-membered rings (Figures 1a and b), and RNA does single-ring bases (Cytosine and Uracil) and double-ring bases (Guanine and Adenine). However, the criteria presented above are appropriate to detect a *π*-*π* stacking: The minimum distance was calculated for all backbone heavy atoms in any ring, and 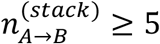 and 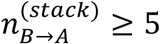 were used for the stacking judgement.

By viewing many snapshots, we found eight stacking patterns when ribocil was in the aptamer’s binding pocket: Three are native stacking and five are non-native stacking (Table S4). Those stacking patterns were found in both the Ribo-A and Ribo-B systems. We denote the numbers of native stacking patterns and non-native stacking patterns in a snapshot as 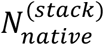 and 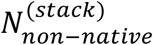, respectively: 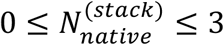 and 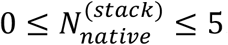.

Figures S7 and S8 are, respectively, the front and side views of some snapshots, which contained the stacking. Figure S7 is presented to show the stacking pattens, and Figure S8 to show the ribocil’s positions relative to X-ray position (PDB: 5c45). These figures are discussed in the main text.

**Table S4.**
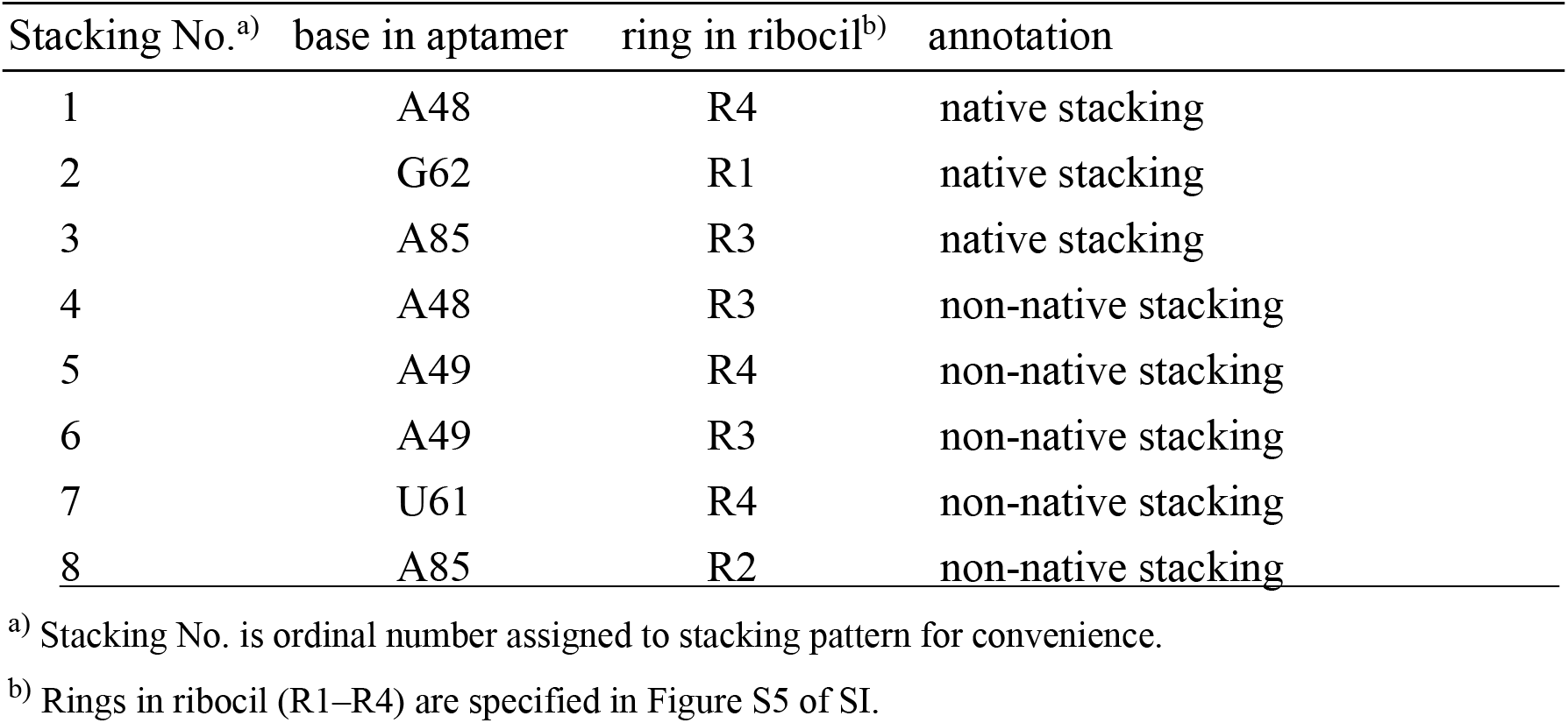
Intermolecular *π*-*π* stacking pairs found in simulation.

| Stacking No.<sup>a)</sup> | base in aptamer | ring in ribocil<sup>b)</sup> | annotation |
| --- | --- | --- | --- |
| 1 | A48 | R4 | native stacking |
| 2 | G62 | R1 | native stacking |
| 3 | A85 | R3 | native stacking |
| 4 | A48 | R3 | non-native stacking |
| 5 | A49 | R4 | non-native stacking |
| 6 | A49 | R3 | non-native stacking |
| 7 | U61 | R4 | non-native stacking |
| 8 | A85 | R2 | non-native stacking |

<sup>a)</sup> Stacking No. is ordinal number assigned to stacking pattern for convenience.
<sup>b)</sup> Rings in ribocil (R1–R4) are specified in Figure S5 of SI.

**Figure S7:**
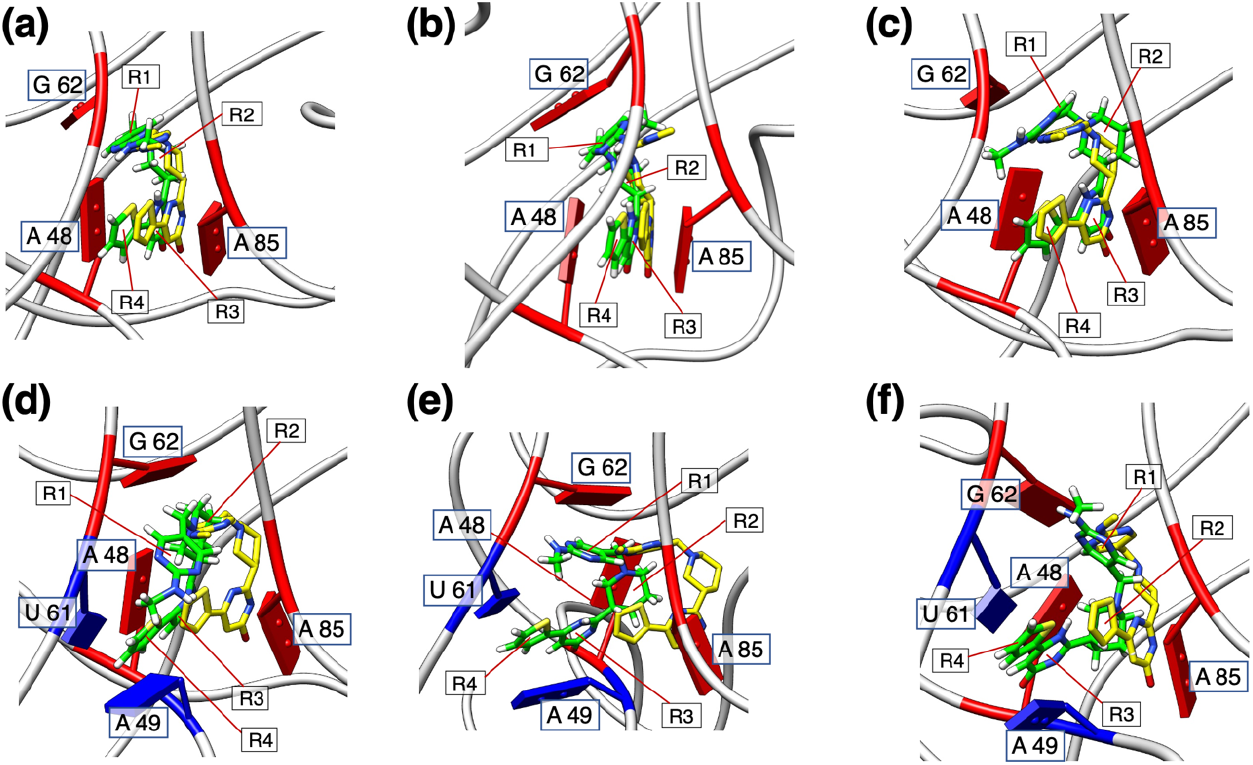
Snapshots with stacking between aptamer domain and ribocil B (green), with focusing ribocil and its surroundings. Light-yellow models are ribocil B in the X-ray complex structure (PDB: 5c45). Panels (a), (b) and (c) are those with native stacking (see Table S4). Panels (d), (e), and (f) are those with non-native stacking: (d) Snapshot with stacking patterns A48–R3, A49–R4 and U61–R4. (e) Snapshot with stacking patterns A49–R4, A49–R3 and U61–R4. (f) Snapshot with stacking patterns A49–R4, A49–R3, U61–R4 and A85–R2. All figures are presented from the front view (Figure S1a).

**Figure S8:**
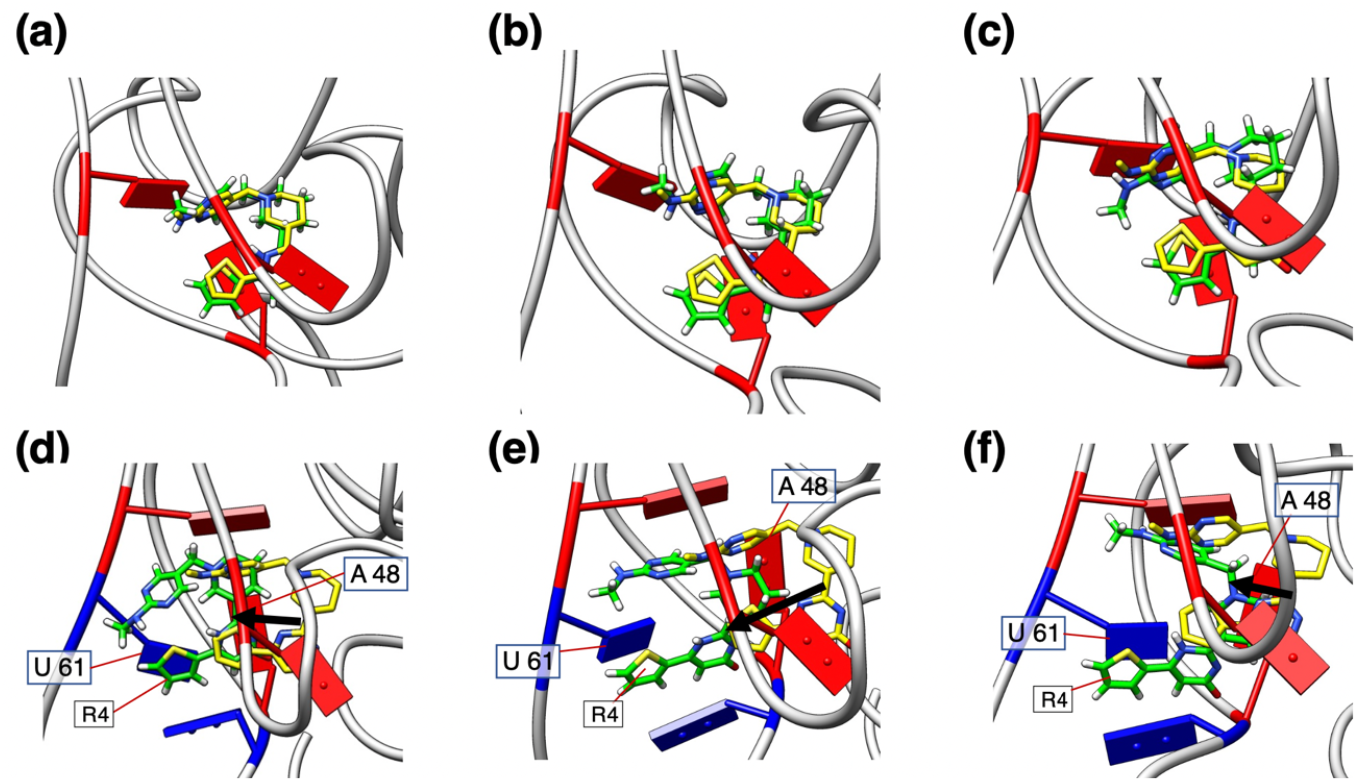
Side view of snapshots in Figure S7. Names of bases of aptamer domain and rings of ribocil, which are shown in Figure S7, are omitted except for A48, U61 and R4. Black arrows in panels (d), (e) and (f) indicate shift of ribocil’s centroid from the X-ray position to the snapshot.

### Section 5. Convergence of *Q*_*cano*_

To access the accuracy of the conformational distribution *Q*_*cano*_(*i, j, k*), which was obtained in the 3D-RC space, we used a function *E*_*local*_(*i, j, k*) (equation 14 of Ref. M1): The smaller the *E*_*local*_ for a zone at a 3D RC index (*i, j, k*), the more accurate the *Q*_*cano*_ in the zone and its adjacent zones. We judged that a region satisfying *E*_*local*_(*i, j, k*) ≤ 0.25 has an appropriate accuracy, where 0.25 is the criteria used repeatedly in our previous studies (Higo et al., 2020b; Hayami et al., 2021; Higo et al., 2021; Higo et al., 2022). Here, we denote this region as *R*_*acc*_(0.25).

We performed 45 iterations as mentioned in the *Results* section of the main text. Figure S9 illustrates the *R*_*acc*_(0.25) region at six iterations together with the iso-density region of *Q*_*cano*_(*i, j, k*) ≥ 1.0 × 10^−5^ (the green-colored contour region). This figure indicates that the *R*_*acc*_(0.25) region increased from iteration 1 to 40, and that the region almost saturated in iterations 40–45 with overlapping to the main body of the iso-density region of *Q*_*cano*_(*i, j, k*) ≥ 1.0 × 10^−5^.

The periphery of the iso-density region was not involved in the *R*_*acc*_(0.25) region, which means that *Q*_*cano*_(*i, j, k*) was inaccurate in the periphery. We, however, note that the density of the periphery region was very small (*Q*_*cano*_(*i, j, k*) ≈ 1.0 × 10^−5^). Therefore, this inaccuracy of *Q*_*cano*_(*i, j, k*) in the periphery region does not cause a serious effect on the main result of the current study.

**Figure S9:**
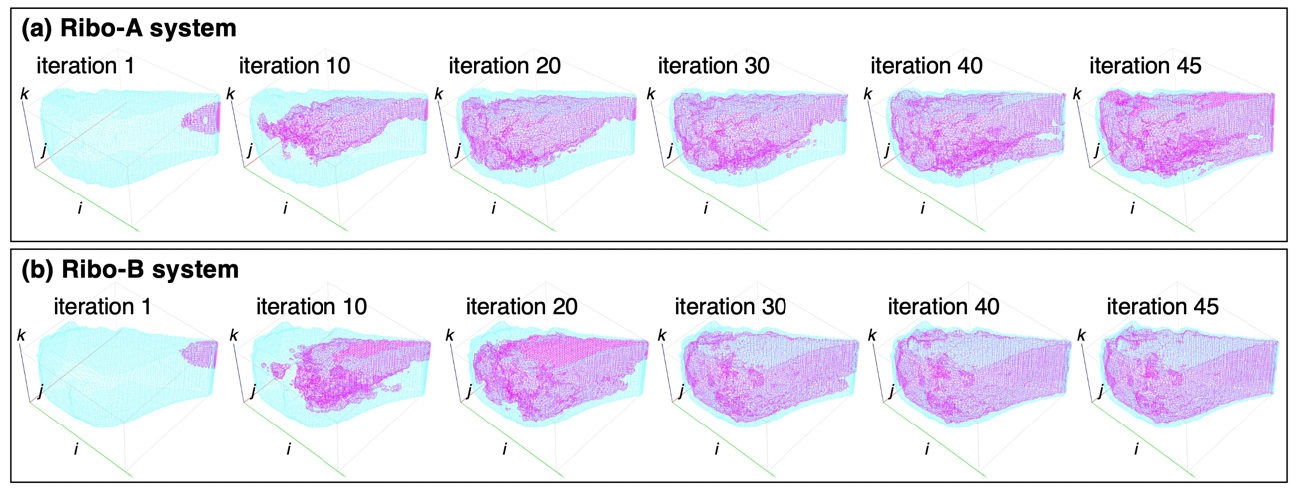
Magenta contours are regions of *R*_*acc*_(0.25) at six iterations for (a) Ribo-A and (b) Ribo-B systems. Iteration No. is shown in the figure. See caption of Figure 2 of main text for the 3D-RC coordinate axes *i, j* and *k*. Cyan contours indicate iso-density region of *Q*_*cano*_(*i, j, k*) = 1.0 × 10^−5^ (Figure 2).

**Figure S10:**
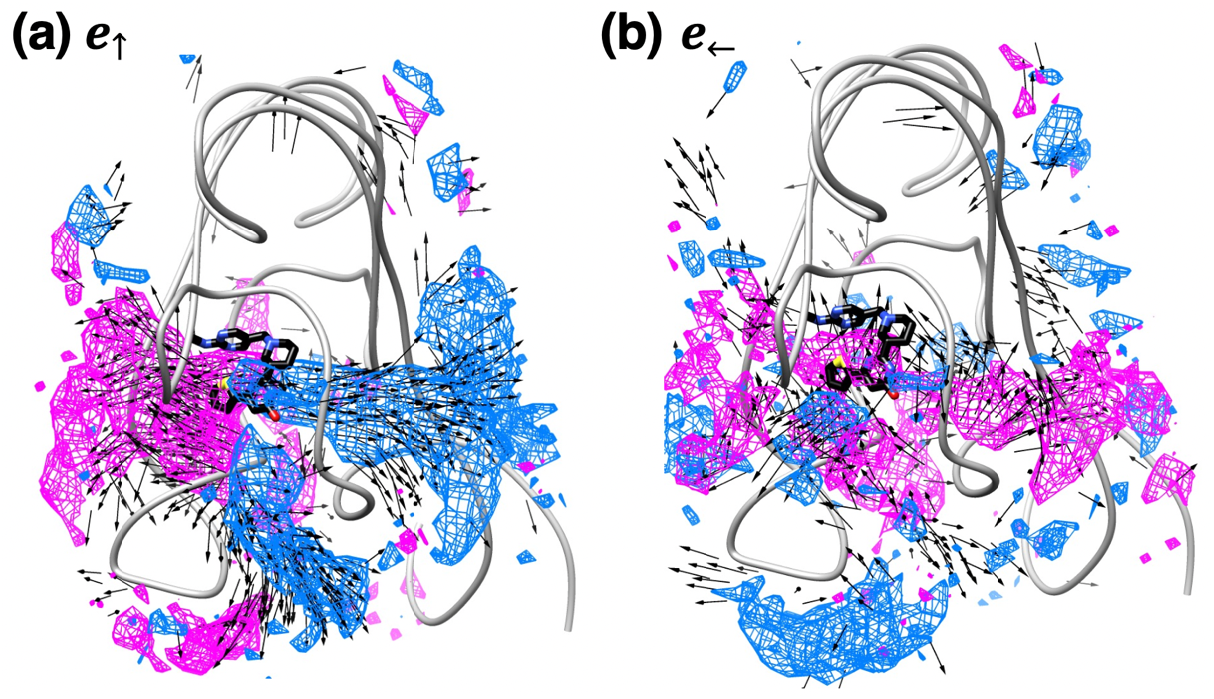
(a) < ***e***_↑_(***r***_*cube*_) > and *SP*_↑_(***r***_*cube*_) for Ribo-A system. (b) < ***e***_←_(***r***_*cube*_) > and *SP*_←_(***r***_*cube*_) for the Ribo-A system. In both panels, vectors < ***e***_*α*_(***r***_*cube*_) > (*α* =↑ or ←) with | < ***e***_*α*_ (***r***_*cube*_) > | ≥ 0.6 are shown by black arrows. Regions with *SP*_*α*_ (***r***_*cube*_) ≥ 0.6 and *SP*_*α*_ (***r***_*cube*_) ≤ −0.6 are shown, respectively, by magenta- and cyan-colored contours. Figure 6 of main text displays < ***e***_*α*_ (***r***_*cube*_) > and *SP*_*α*_ (***r***_*cube*_) for the Ribo-B system.

**Figure S11:**
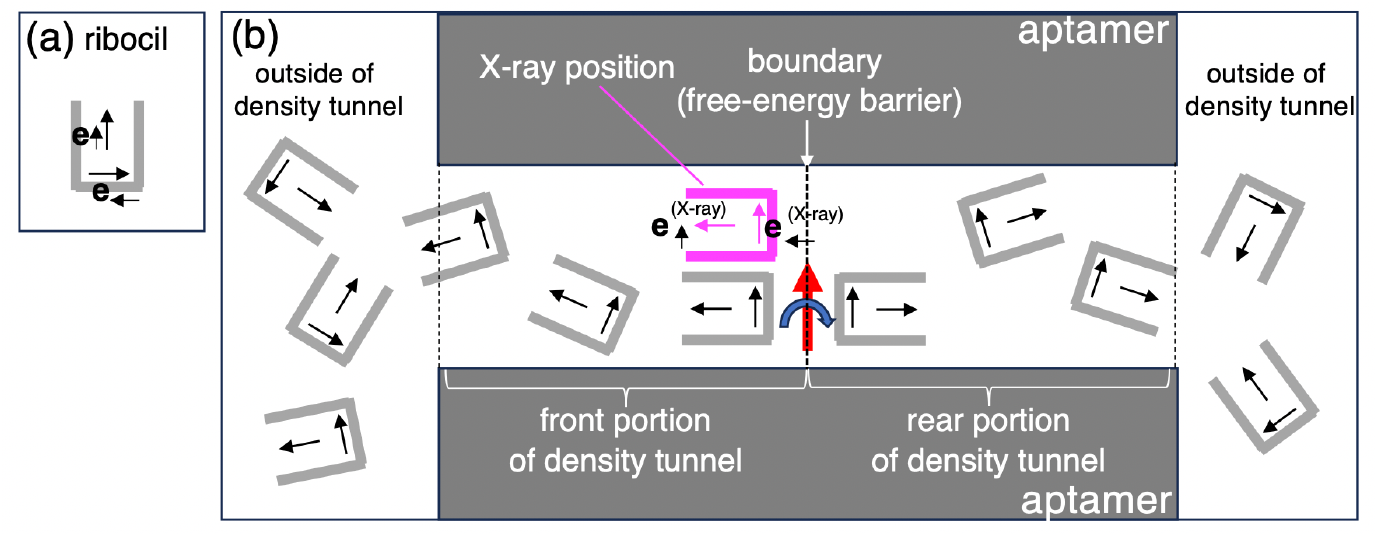
(a) Orientation vectors ***e***_↑_ and ***e***_←_ set to ribocil. See Subsection 4.2. of SI doe details. (b) Behavior of ***e***_↑_ and ***e***_←_ outside and inside the density tunnel. Orientation vectors are randomized outside the tunnel. When ribocil is in the front portion of the tunnel, ***e***_↑_ tends to be parallel to 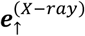, and ***e***_←_ does to be parallel to 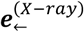 weakly. 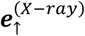 and 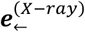 are respectively ***e***_↑_ and ***e***_←_ of ribocil in the X-ray position (magenta ribocil). When ribocil is in the rear portion of the tunnel, ***e***_↑_ tends to be anti-parallel to 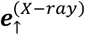, although ***e***_←_ tends to be parallel to 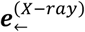 weakly. Because < ***e***_↑_ > varies sharply at the boundary of the front and rear portions of the density tunnel, ***e***_↑_ rotates around the rotation axis (red arrow) located at the boundary. This large rotation of ***e***_↑_ induces a free-energy barrier at the boundary, and the barrier is visualized in the spatial patterns of *ρ*_*CM*_ (broken line of Figure 5).

**Figure S12:**
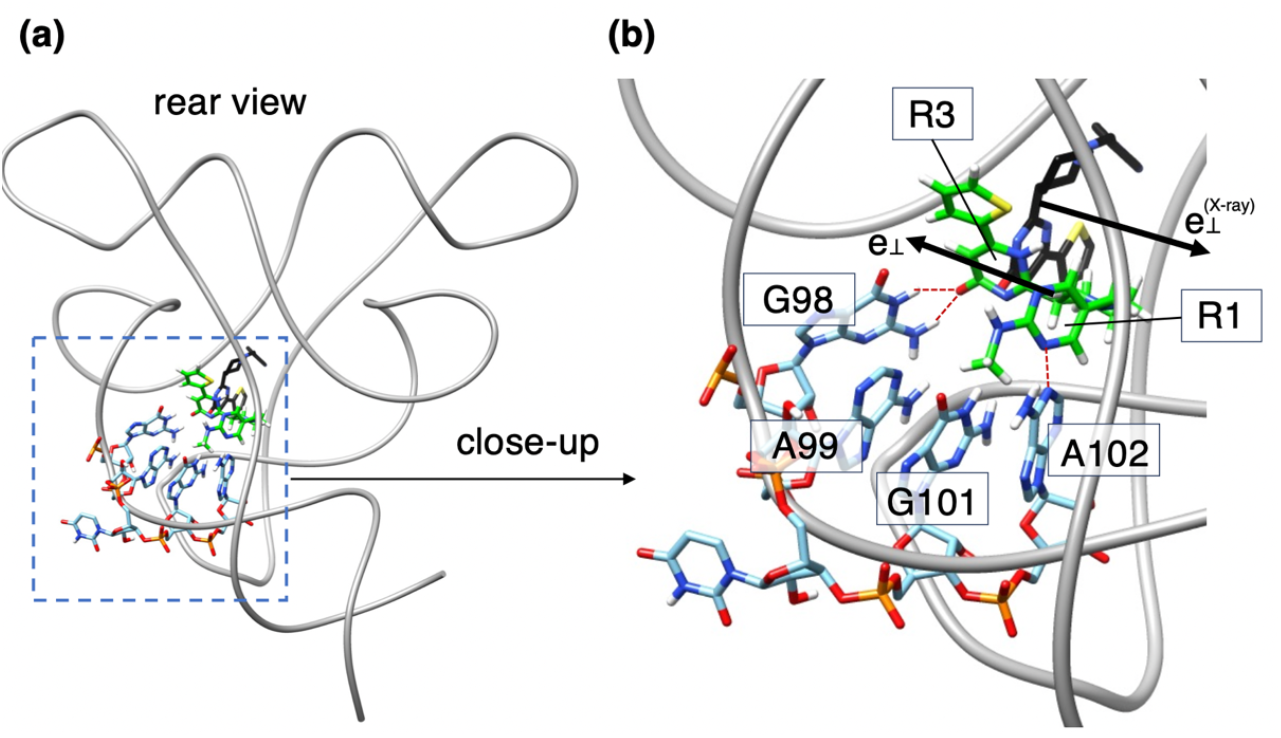
(a) Rear view of a snapshot, where ribocil B (green) is in rear portion of density tunnel. Ribocil in black is that in the X-ray complex structure (PDB: 5c45). (b) Close-up of broken-line rectangle in panel (a). Aptamer domain is illustrated in two models: Entire aptamer domain is presented by ribbon model, and bases 98–102 are by stick model. Brown broken lines are hydrogen bonds between ribocil and aptamer domain. ***e***_↑_ and 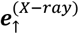 are shown by black arrows, whose definition is shown in Subsection 4.2. of SI. Two rings, R1 and R4, of ribocil are defined in Figure S5.

